# Evolution of Natural Lifespan Variation and Molecular Strategies of Extended Lifespan

**DOI:** 10.1101/2020.11.09.374488

**Authors:** Alaattin Kaya, Cheryl Zi Jin Phua, Mitchell Lee, Lu Wang, Alexander Tyshkovskiy, Siming Ma, Benjamin Barre, Weiqiang Liu, Benjamin R. Harrison, Xiaqing Zhao, Xuming Zhou, Brian M. Wasko, Theo K. Bammler, Daniel E. Promislow, Matt Kaeberlein, Vadim N. Gladyshev

## Abstract

To understand the genetic basis and selective forces acting on longevity, it is useful to examine lifespan variation among closely related species, or ecologically diverse isolates of the same species, within a controlled environment. In particular, this approach may lead to understanding mechanisms underlying natural variation in lifespan. Here, we analyzed 76 ecologically diverse wild yeast isolates and discovered a wide diversity of replicative lifespan. Phylogenetic analyses pointed to genes and environmental factors that strongly interact to modulate the observed aging patterns. We then identified genetic networks causally associated with natural variation in replicative lifespan across wild yeast isolates, as well as genes, metabolites and pathways, many of which have never been associated with yeast lifespan in laboratory settings. In addition, a combined analysis of lifespan-associated metabolic and transcriptomic changes revealed unique adaptations to interconnected amino acid biosynthesis, glutamate metabolism and mitochondrial function in long-lived strains. Overall, our multi-omic and lifespan analyses across diverse isolates of the same species shows how gene-environment interactions shape cellular processes involved in phenotypic variation such as lifespan.

## Introduction

Diverse selective forces (mutation, selection, drift) generate enormous variation within and among species. [1, 2]. Consequently, many morphological, behavioral and physiological phenotypes (traits) vary within and between species in natural populations [3–7]. For example, genetic variation in natural populations of many organisms can differentially affect their neural and endocrine functions, leading to variation in quantitative life-history traits such as fitness and age at maturation [8, 9]. Variation in another fitness trait, lifespan, has also attracted much attention [10–12]. Across eukaryotic species, longevity can differ over many orders of magnitude, from days to centuries, e.g. the Greenland shark (*Somniosus microcephalus*) may live for more than 500 years whereas some species live only several days [13–15]. Longevity also varies among individuals of the same species, indicating that variability of lifespan is not constrained at the level of species, and that the molecular determinants of lifespan vary within the same genetic pool [16–21].

What, then, are the factors that determine the lifespan of individuals? The molecular pathways that underlie the genetic and environmental determinants of lifespan are among the most intensely studied areas in the aging field [18, 19, 22–26]. Laboratory animal models have shown that longevity can be extended by environmental [27–30], dietary [31–33], pharmacological [34–37] and genetic interventions [38–40]. However, many of these laboratory-adapted populations are constrained by genetic and environmental background [41–43]. For example, artificially created mutant strains may show longer lifespan under laboratory settings but demonstrate reduced fitness in their natural environment [44–47]. A more integrated approach is needed to understand how the natural environment and natural selection interact to shape lifespan and associated life-history traits.

In order to better understand the impact of natural genetic variation on lifespan, we studied 76 wild isolates of the budding yeast to capture the molecular signatures of evolved diversity of lifespan. This collection included 40 diploid isolates of *Saccharomyces cerevisiae* and 36 diploid isolates of *Saccharomyces paradoxus* [48, 49]. Their niches include human-associated environments, such as breweries and bakeries, and different types of wild ecological niches, such as trees, fruits, vineyards, and soils across different continents. There was also a group of clinical isolates; *S. cerevisiae* strains isolated from immunocompromised patients. *S. cerevisiae* and *S. paradoxus* are closely related (share a common ancestor between 0.4 and 3 million years ago), with 90% genome identity, and can mate and produce viable progeny [50, 51]. Earlier genome sequencing of these isolates revealed allelic profiles, their ploidy status, and a phylogeny of these isolates [48–50]. Based on these studies, it was shown that while some of these strains fall into distinct lineages with unique genetic variants, almost half of the strains have mosaic recombinant genomes arising from outcrosses between genetically distinct lineages of the same species [49]. Because of their wide ecological, geographical, and genetic diversity, natural isolates of the budding yeast *Saccharomyces* have become an important model system for population/evolutionary genomics [52] and to study the complex genetic architecture of lifespan [18, 53–57]. Therefore, these strains offer a powerful genetic pool to understand how natural genetic variation may shape lifespan variation.

Accordingly, we assayed the replicative lifespan (RLS) of ~3,000 individual cells representing these isolates under two different conditions: yeast peptone dextrose (YPD, with 2% glucose), and yeast peptone glycerol (YPG, 3% glycerol as a respiratory carbon source), and identified up to 10-fold variation in median RLS under each condition. Although little is known about the life histories of these wild isolates they face different, niche-specific evolutionary pressures for adaptation to different stresses [49–52]. To understand how different genotypes arrive at different lifespan phenotypes, we further analyzed endophenotypes (gene expression, metabolite abundance) to characterize the molecular patterns associated with condition-specific lifespan variation. Following characterization of transcripts and metabolites with significant association with longevity, we identified pathways associated with median RLS across these isolates. Our data showed that the naturally arising variation in genotype can cause large differences in lifespan, which are associated with distinct patterns of gene expression and metabolite abundances. These analyses revealed connected pathways that have not been previously associated with lifespan variation in a laboratory setting. Altogether, we present the most comprehensive analysis to date of how the environment and genetic variation interact to shape aging and the associated life-history traits.

## Results

### Growth characteristics and lifespan variation across wild isolates

First, we monitored growth characteristics of natural yeast isolates on standard glucose media (i.e. during fermentation) and on media with a non-fermentable substrate, glycerol, as a carbon source (i.e. during respiration), using an automated growth analyzer, and calculated the doubling time. Most wild isolates grew faster than the diploid laboratory wild type (WT) BY4743 strain under both conditions, with an average doubling time of 65 minutes in YPD and 125 minutes in YPG **(Fig. 1A, Table S1)**. Importantly, most of these strains grew at a similar rate on YPD, and variation in growth rate on YPG was also relatively small among them, indicating that these laboratory-optimized culture conditions are suitable for supporting nutritional needs of these strains and for RLS analysis **(Fig. 1A)**.

**Figure 1:**
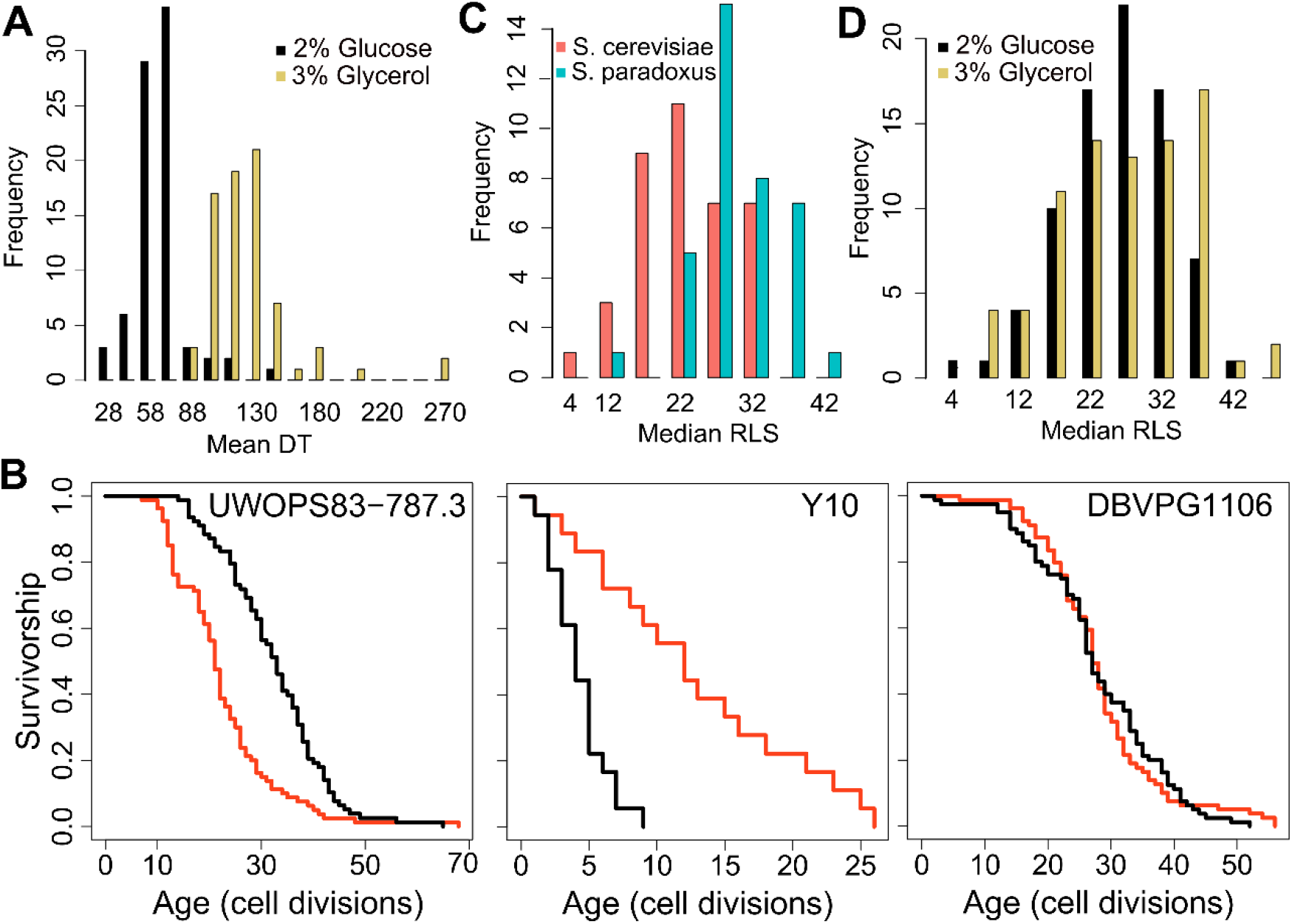
Doubling time and replicative lifespan of yeast wild isolates. **A)** Distribution of mean doubling time (DT, in minutes) on YPD (2% glucose) and YPG (3% glycerol). **B)** Examples of lifespan curves for the selected strains. Black curve shows lifespan under YPD conditions, and red curve under YPG conditions. **C)** Median replicative lifespan (RLS) distribution across *S. cerevisiae* (red) and *S. paradoxus* (turquoise) isolates grown in YPD. **D)** Distribution of median RLS across different conditions. Source data is provided as Supplementary Table 1.

Next, we assayed RLS of these isolates at 30°C on both growth conditions (YPD and YPG media). Under the YPD condition, we observed a remarkable ~10-fold variation in median and maximum RLS (Pearson correlation coefficient = 0.65 between median and maximum RLS, p < 0.00001) across these isolates **(Fig. 1B, 1C, Fig. S1A, Table S1)**. The average median RLS of *S. paradoxus* strains (29.7) was significantly higher (p = 2.7×10^−9^) than the average median RLS of *S. cerevisiae* strains (22.7) (**Fig. 1C, S1A**). Among the 76 strains analyzed, *S. paradoxus* strain Y7 showed the longest median RLS (RLS=42), whereas *S. cerevisiae* strain Y10 exhibited the shortest median RLS, with 50% of Y10 cells ceasing division after producing only four daughters **(Table S1)**.

Glycerol as a growth substrate can extend both RLS and chronological lifespan (CLS) [58, 59]. In the case of CLS, the increased longevity is caused by a switch from fermentation to respiration [58]; however, mechanisms by which glycerol affects RLS are unclear, since respiratory metabolism is not always required for RLS extension in laboratory WT strains [59]. While we observed a significant (p < 0.05, Wilcoxon Rank Sum Test) median RLS increase in 32 strains on YPG with 24 strains significantly decreased median RLS; and the remaining seven strains showed no significant changes (**Fig. 1D, Fig. S1B, Table S1**). For example, *S. cerevisiae* strain BC187 showed a significantly increased median RLS on glycerol (37.5 in YPG versus 32 in YPD), and. *S. paradoxus* strain KPN3829 also showed a similar increase (21 in YPD and 35 in YPG) **(Table S1)**. On the other hand, *S. cerevisiae* strain YJM975 showed a significant decrease (median RLS = 31 in YPD and 23 in YPG) **(Table S1)**.

We further dissected the effect of carbon source on RLS by comparing median RLS variation between YPD and YPG conditions. Interestingly, strains with the shortest RLS on YPD tended to achieve the longest lifespans on YPG, while the long-lived strains on YPD generally did not show a further RLS increase (**Fig. S1B, S1C** Pearson correlation coefficient = −0.51 between median RLS on YPD and median RLS on YPG, p < 0.0001). A similar observation was shown under CR conditions in the case of single-gene deletions, where the shortest-lived strains tended to yield the largest lifespan extension when subjected to CR [60]. Overall, we observed significant differences in lifespan across these strains on both conditions. The observed lifespan pattern on YPG conditions in comparison to those on YPD suggests that long-lived strains might reside in environments with low fermentable carbon sources, so that they undergo distinct metabolic regulation to metabolize respiratory carbon sources. As such, they did not show further RLS extension when grown in glycerol.

### Endophenotype variation across wild isolates

While many previous large-scale omics studies on aging have focused on genome-wide association [53, 61–63], recent comparative studies on transcriptomics [64, 65], proteomics [66–68], metabolomics [69–71] and ionomics [72] have begun to shed light on molecular patterns and mechanisms associated with the aging process. For example, it has been suggested that natural variation is associated with extensive changes in gene expression, translation and metabolic regulation, which in turn may affect fitness under different stress conditions [73, 74]. In fact, gene expression variation has repeatedly been postulated to play a major role in adaptive evolution and phenotypic plasticity [74, 75], as well as specific phenotypic outcomes such as changes in morphology [76] and lifespan [77]. Similarly, comparative studies of metabolite profiles have been utilized to describe the genotype to phenotype relations in model organisms [78, 79]. Accordingly, we aimed to explain lifespan differences among these wild-derived yeast isolates by analyzing their gene expression variation (based on transcriptomics analyses) and differences in their metabolite levels (based on metabolomics analyses).

In the case of the transcriptome, we obtained ≥ 5 million 150-bp paired-end RNA-seq reads for each strain grown on YPD. For metabolomics analyses, we applied targeted metabolite profiling using liquid chromatography-mass spectrometry (LC-MS). After filtering and quality control, the data set contained RNA-seq reads for 5,376 genes and 166 metabolites identified commonly across all isolates (**Table S2**). The expression profiles between strains were similar to one another, with Spearman correlation coefficients of strain pairs ranging between 0.59 and 0.93 (except the pairing involving Q59.1, CBS5829, YPS606, and UFRJ50791, with the range between 0.21 and 0.79) **(Fig. S2)**. To determine whether the previously published sequence-based evolutionary relationships [49] were reflected in their gene expression variation, we constructed gene expression phylograms using a distance matrix of 1 minus Spearman correlation coefficients and the neighbor-joining method [80]. The resulting topology of species-specific trends was largely consistent with their phylogeny with a clear separation between *S. cerevisiae* and *S. paradoxus* strains **(Fig. 2A)**; however, at the intra-species level, the topology was not consistent with the phylograms of genomic data.

**Figure 2:**
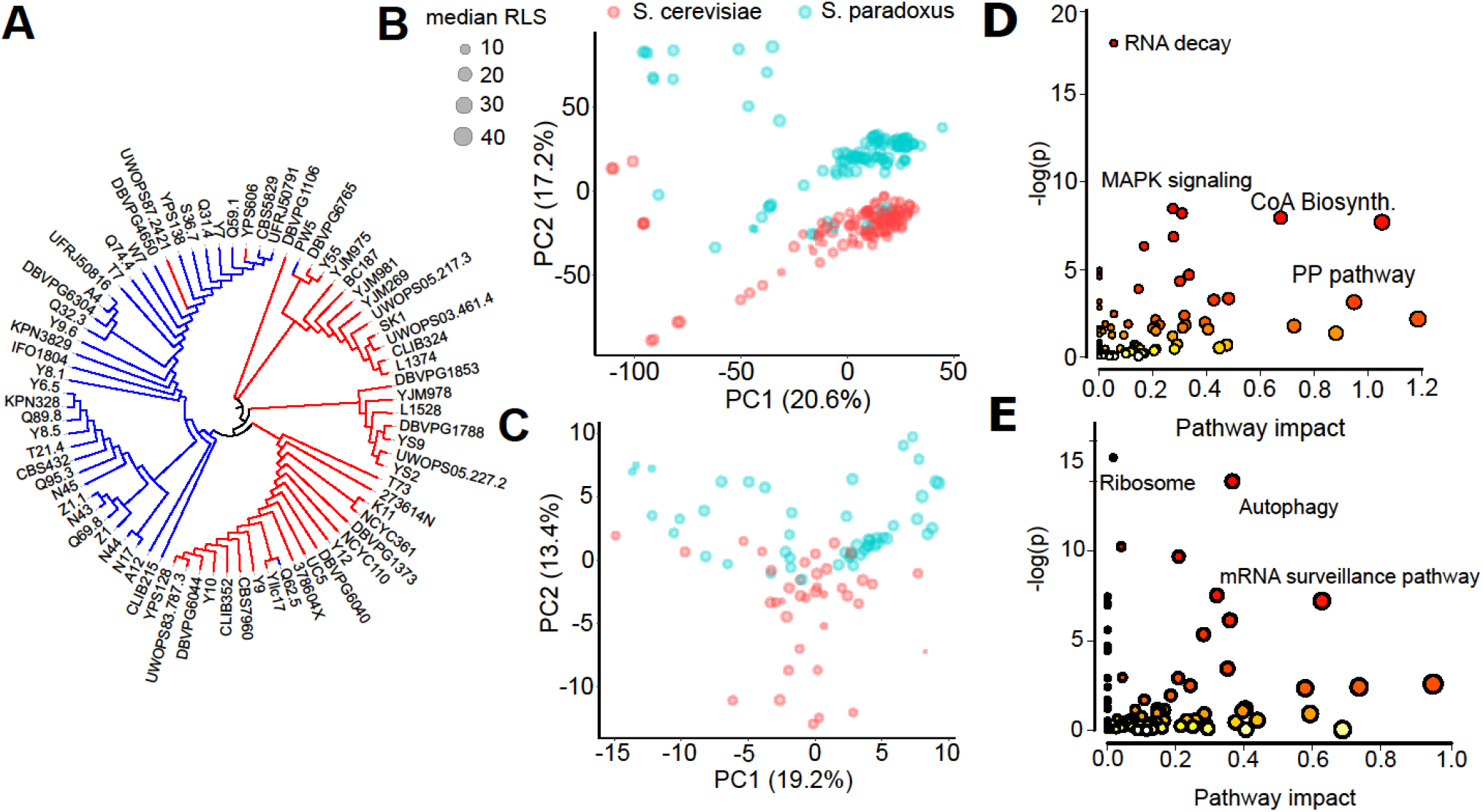
Endophenotypic variation across strains. **(A)** Phylogenetic relationship based on the transcriptome data of 76 strains of two species. Principal component analysis (PCA) of **(B)** transcriptomics and **(C)** metabolomics. Percent variance explained by each principal component (PC) is shown in parentheses. Pathway enrichment analysis for combined top genes and metabolites contributing to **(D)** PC1 and **(E)** PC2. Some of the enriched KEGG pathways are shown in each panel. PCA loadings can be found in Supplementary Table 3.

To visualize endophenotypic variation between these two species and across the strains of the same species, we performed principal component analysis (PCA) on each type of data. PCA of the transcriptome revealed a pattern resembling the phylogenetic relationship, with the first three PCs explaining ~49 % of total variance in gene expression **(Fig. 2B, Fig. S3A)**. Although some *S. paradoxus* strains clustered with *S. cerevisiae*, we observed clear species segregation based on PC2 (except for some outlier strains that were separated by PC1). PCA of metabolomics data revealed a similar structure with the first three PCs explaining ~41% of total variance in metabolite levels, and PC2 somewhat separating the species **(Fig. 2C, Fig. S3B)**.

To understand the basis of this segregation pattern, we performed pathway enrichment analysis by combining the 500 top genes (250 with positive weights and 250 with negative weights) and 40 top metabolites (20 with positive weights and 20 with negative weights) contributing to each PC, respectively. This integrative analysis of genes and metabolites (see Materials and Methods) contributing to PC1 revealed a distinct set of Kyoto Encyclopedia of Genes and Genomes (KEGG) pathways, including RNA degradation, MAPK signaling pathway, cell cycle, pantothenate and CoA biosynthesis, ribosome biogenesis and pentose phosphate pathway **(Fig. 2D, Table S3)**. The analysis of genes and metabolites for PC2 revealed the KEGG pathways related to ribosome, autophagy, endocytosis, cell cycle, mRNA surveillance, and nucleotide excision repair **(Fig. 2E, Table S3)**. These results suggest that these processes diverged most significantly across the wild isolates of two species of *Saccharomyces* genus and may account for their phenotypic diversity, including lifespan. It should be noted, however, that we do not know exactly what each PC represents, unless it perfectly aligns or correlates with some known variables. In addition, either biological (e.g. phylogenetic structure), technical (e.g. data normalization or batch effect), or mixed effects of both may render PCA biased [81].

### Relationship between endophenotypes and lifespan

To identify endophenotypes (transcripts and metabolites) correlating with lifespan variation across wild isolates, we applied the phylogenetic generalized least-squares (PGLS) method to account for phylogenetic relationships among the strains and test for different models of trait evolution [82, 83]. Regression was performed between endophenotypic values and median RLS under different models of trait evolution and the best-fit model was then selected based on maximal likelihood. To assess the robustness of these relationships, we repeated the regression after taking out one yeast strain at a time and only those regressions that remained significant were further considered. This ensured the overall relationship did not depend on a particular isolate.

With the PGLS approach, we identified 73 transcripts with significant correlation with median RLS (*Padj ≤ 0.01*; 39 with positive correlation and 34 with negative correlation) **(Table S2)**. Among the top hits with positive correlation were a putative zinc finger protein coding gene *CMR3* (*Padj*=3.3×10^−9^), histone acetyltransferase (HAT) gene *HPA2* (*Padj*=0.0002), Transcription factor *TEC1* (*Padj*=9.3×10^−6^) and zing regulated protein gene *ZRG8* (*Padj*=0.006) **(Fig. 3A, Table S2)**. The top hits with negative correlation included the genes coding for cyclin-dependent kinase Pho85p interacting proteins *PCL1* (*Padj*=0.0008) and *PCL2* (*Padj*=0.001), regulator of Ty1 transposon protein coding gene *RTT107* (*Padj*=0.007), and inositol monophosphatase gene *INM1* (*Padj*=0.006) **(Fig. 3B, Table S2)**. Next, to assess if any of our transcript hits were previously implicated in yeast lifespan, we extended our list of significant genes to 357 genes by selecting a cutoff at *Padj*=0.05 and compared these with the genes associated with RLS in laboratory WT strain listed in the GenAge database [84]. GenAge identifies 611 genes from the published literature with effects on RLS (decreased or increased) of laboratory yeast strains (595 deletion mutants and 16 overexpressed genes) **(Table S4)**. Of 5,376 genes whose expression was measured across the wild isolates, there were 39 genes present in both our list and GenAge, 23 of which showed the same direction of correlation with RLS. For example, *INM1, RTT107, PPH3* and *BSC1* genes increase RLS when deleted (GenAge database) and are associated with increased RLS when their transcript levels decrease across wild isolates (this study). **(Table S4)**. However, the overall pattern of overlapping genes as well as the direction of correlation did not reach statistical significance (Fisher’s exact test, p ≥ 0.05). It should be noted that many of the RLS associated genes listed in GeneAge are reported from single gene KO studies and there has been no comprehensive studies examining gene overexpression on a genome-wide scale. This raises a possibility that the genes we identified here might not necessarily be over-represented among the lifespan-related genes from other studies. It is also possible that the genetic architecture of trait variation in natural populations may differ from that which is assumed from studies of lab strains, including extensive single-gene studies of lifespan variation in yeast [85]. The lack of overlap between the genes whose expression correlates with lifespan variation in wild isolates and genes that affect RLS in single-mutant studies on laboratory WT background supports this possibility. Considering this, we then asked if trait variation in wild isolates and lab strains may converge at the transcriptome in a way that may be detectable at the level of gene expression, or at the level of biological pathway. To do this we examined the gene expression patterns across wild isolates with those of 1,376 laboratory knock-out strains (KO) strains [86] whose RLS was quantified [85] previously **(Fig. 4, Fig. S4A, S4B Table S1)**. We calculated an association of gene expression with different measures of RLS (mean RLS, median RLS, and maximum RLS) across KO strains. Our analysis revealed around 400 significant genes (*Padj*=0.05) associated with three types of RLS measures, and more than 1000 genes associated with median RLS **(Fig. 4A)**. To compare the RLS-associated transcriptomes of wild isolates and lab strains, we then calculated a correlation matrix of RLS-associated gene expression changes across KO strains and wild isolates **(Fig. 4B)**. We found no positive correlation between RLS associated gene expression changes across KO strains and RLS associated gene expression changes across natural isolates **(Fig. 4B)**.

**Figure 3:**
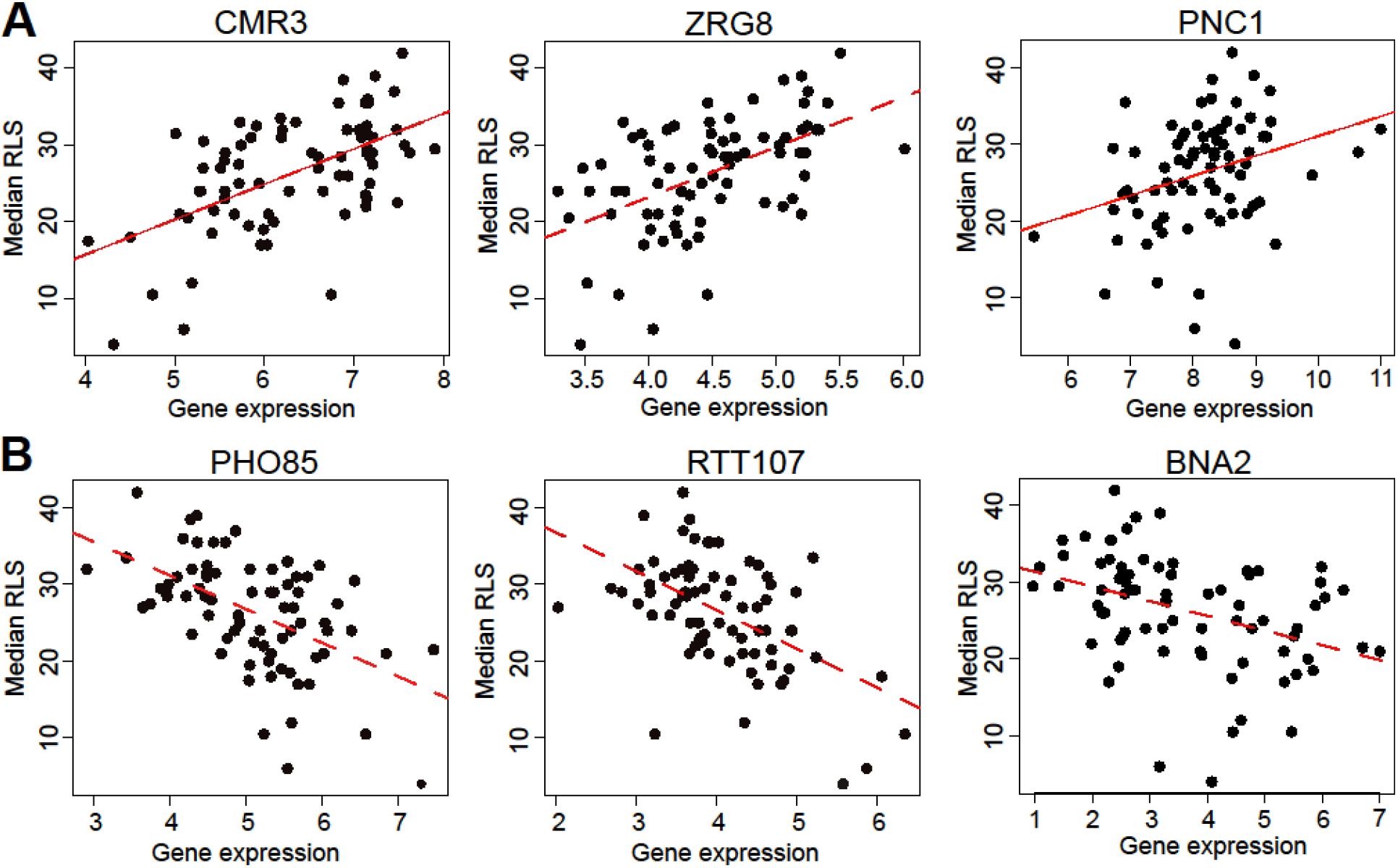
Selected genes whose expression correlates with median RLS. **(A)** Gene expression level (log2-cpm) of *CMR3, ZRG8*, and *PNC1* positively correlates with median RLS. **(B)** Transcript abundance of *PHO85, RTT107*, and *BNA2* negatively correlates with median RLS. Regression slope *P* values can be found in Supplementary Table 2, which is also the source data file for these analyses.

**Figure 4:**
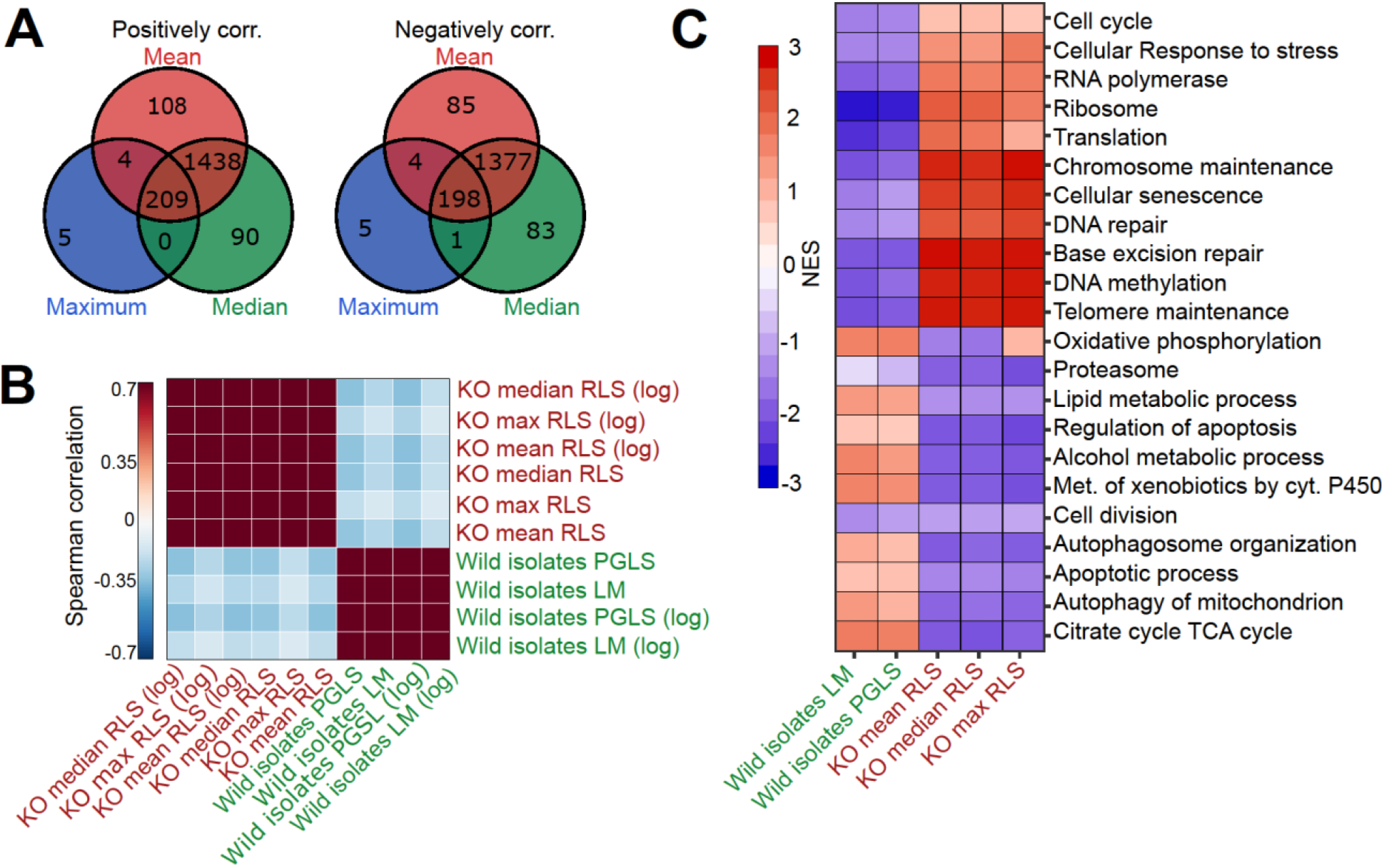
Comparative analysis of lab yeast knockout (KO) and wild isolates. **(A)** Significant genes (*Padj* < 0.05) associated with maximum, median and mean RLS (or log maximum, median and mean) across deletion strains based on transcriptomics data obtained from 1,376 KO strains. Genes positively and negatively associated with RLS (upregulated and downregulated, respectively) are significantly shared across different metrics of RLS (Fisher exact test p < 0.05). **(B)** Denoised correlation matrix of gene expression effects across single-gene deletion strains (KO), and those that we measure across the wild isolates that are associated with RLS. Correlation coefficient is calculated using union of top 1,000 statistically significant genes for each pair of signatures with Spearman method. LM: Linear model; PGLS: phylogenetic regression least squares. **(C)** Functional enrichment of genes associated with RLS across deletion and natural strains. Cells are colored based on normalized enrichment score (NES). Supplementary Table 1 and 2 are provided as source data files for these analyses.

We then performed functional enrichment (GSEA) of genes associated with RLS across deletion and wild isolates to see if associations with RLS may converge at the level of the biological pathway. We find that the transcripts associated with RLS in these two populations enrich distinct sets of biological pathways **(Fig. 4C)**. For the genes correlating positively with longevity across the KO strains, the enriched terms included cellular responses to stress, ribosome, translation, cellular senescence, and DNA repair **(Fig. 4C)**. On the other hand, terms enriched in wild isolates included TCA cycle, oxidative phosphorylation, and lipid metabolic process, regulation of apoptosis, and autophagy **(Fig. 4C)**. Overall, our comparative analyses of lifespan associated gene expression signatures in laboratory adapted yeast strains versus wild isolates suggest that different genetic trajectories might have evolved at transcript level across wild isolates to regulate lifespan.

Next, we searched for metabolites whose abundances associate with RLS across wild yeast isolates. The metabolome represents a snapshot of regulation downstream of both the transcriptome, and proteome and it has been effectively used for characterizing phenotypic variation that includes lifespan [69–71]. Among 166 metabolites that we examined, 31 exhibited significant association with median RLS (*Padj ≤ 0.05*) **(Table S2)**. Among the top hits, tryptophan, lactate, 2-hydroxyglutarate, 3-hydroxypropionic acid, 2-hydroxyisobutyrate, 2-hydroxybutyrate, and phenyllactic acid correlated positively **(Fig. 5A)**, whereas lysine, quinolinic acid, propionate, Se-methylselenocysteine showed negative correlation **(Fig. 5B)**. Our metabolite list also included several related short chain fatty acids, with positive correlation to RLS (SCFAs: 3-hydroxypropionic acid, 2-hydroxyisobutyrate, 2-hydroxybutyrate and 2-hydroxyglutarate), which are known to be involved in redox regulations, epigenetic modification, and energy generation [87–88]. Having identified transcripts and metabolites associated with lifespan, we aimed to investigate interaction among them to better understand biological causes of lifespan variation and mechanisms of longevity across the wild isolates.

**Figure 5:**
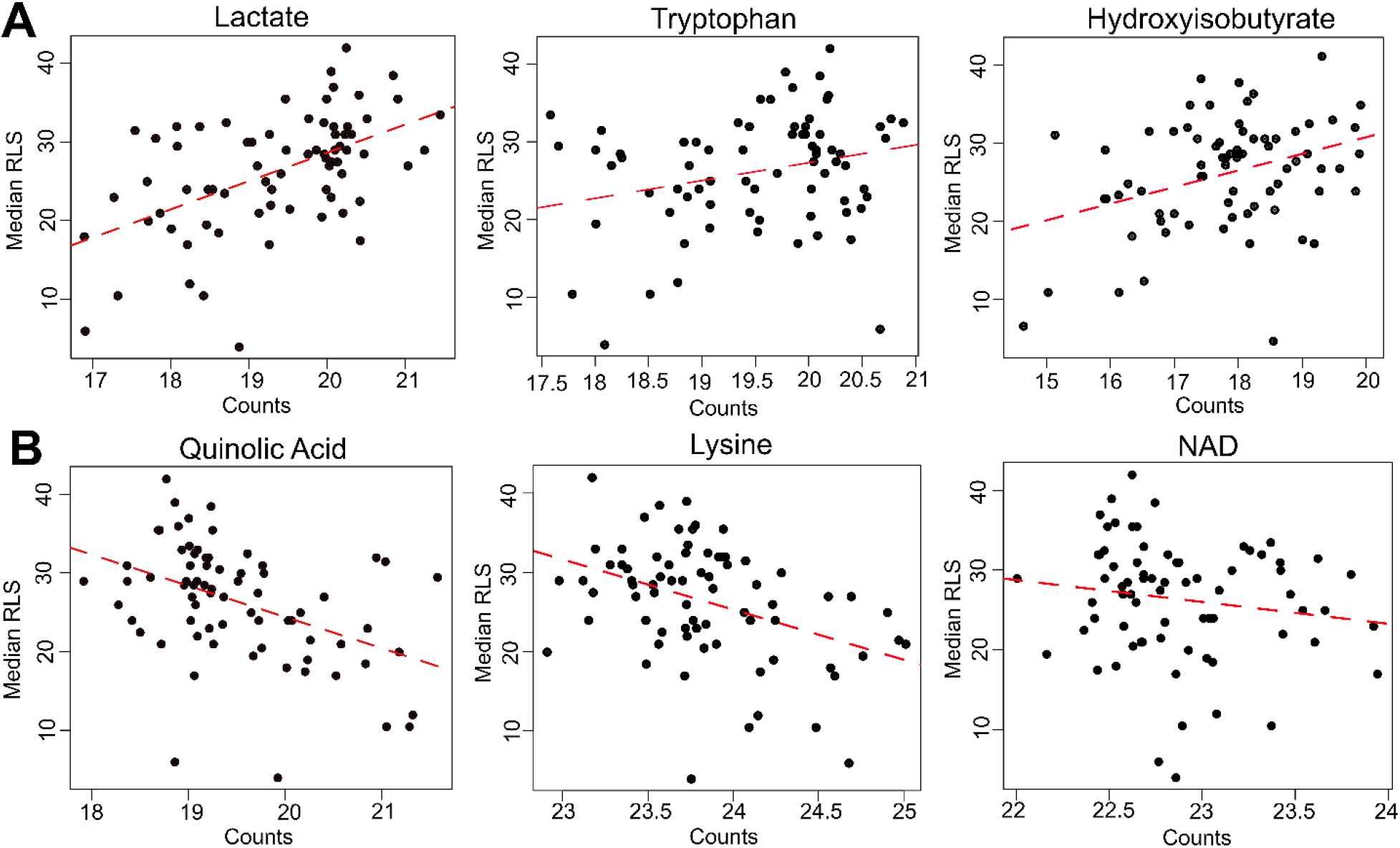
Selected metabolites correlating with median RLS. **(A)** Abundance (LC-MS counts) of lactate, Trp, and hydroxyisobutyrate that positively correlate, and **(B)** abundance of quinolinic acid, Lys, and NAD that negatively correlate with median RLS. Regression slope *P* values can be found in Supplementary Table 2. This file is also provided as a source data file for these analyses.

### Molecular signatures of lifespan extension across wild isolates

To understand the molecular basis of RLS variation in wild isolates, we applied an integrated pathway approach [89]. A combined metabolomics and transcriptomics data analysis revealed a potential role for differential metabolic regulation of tryptophan (Trp), lysine (Lys) and branched chain amino acid (BCAAs) biosynthesis as well as valine (Val) and isoleucine (Iso). For example, our metabolome data revealed that Trp abundance correlates positively with RLS (**Fig. 5**). In addition, we found that quinolinic acid (QA), an intermediate in the Trp catabolic pathway (also known as the kynurenine (KYN) pathway) [90, 91] correlates negatively with median RLS (**Fig. 5**), suggesting a possible inhibition of Trp degradation, corresponding to an increased Trp abundance in long-lived strains. Further evidence for metabolic regulation of Trp in long-lived strains came from our transcriptome data wherein *BNA2* (indoleamine 2,3-dioxygenase) gene, which supports the first rate-limiting step of Trp catabolism [91], was significantly downregulated (*Padj* = 0.02) in long-lived wild isolates **(Fig. 3B, Fig. S5)**. These observations draw a complete picture for the observed Trp abundance in long-lived strains.

Similarly, a link between our transcriptome and metabolome data provided insights into Lys metabolism. We observed a negative correlation between Lys abundance and lifespan (i.e., long-lived strains tend to have less Lys) (**Fig. 5**). Additionally, our transcriptome data showed negative correlations of two homocitrate synthase genes, *LYS20* and *LYS21*, controlling the first rate-limiting step of Lys biosynthesis by catalyzing condensation of Acetyl-CoA and alpha-ketoglutarate (**α**-KG) to produce homocitrate **(Fig. S6, Fig. S7)**. Together, these observations support the idea of decreased Lys levels in long-lived strains. It is also of interest that while they did not reach significance, all genes (with the exception of *ARO8*) involved in Lys biosynthesis showed a trend for decreased expression in long-lived strains (**Table S2**). Interestingly, previous studies have shown the connection between Trp and Lys metabolism both at genetic and metabolic levels. For example, it has been shown that 3-hydroxyanthranilic acid, an intermediate from Trp degradation, can be used as a substrate to synthesize α-ketoadipate [92, 93], which is then converted by *ARO8* to Lys (**Fig. S7**). In this regard, glutamate (Glu) dependent *ARO8* activity is involved in both Trp and Lys catabolic pathways (**Fig. S5, S7**). In addition, at the genetic level, while individual knock-out lines of *BNA2, LYS20* or *LYS21* are not lethal, it was found that the combined deletion of *BNA2* and *LYS20 or BNA2* with *LYS21* causes synthetic lethality [94], possibly by causing Lys auxotrophy. In the light of these observations, our data suggest that the observed occurred differences in Trp and Lys metabolism are not random. Decreased Trp catabolism in long-lived strains might limit α-ketoadipate production, which in turn could affect Lys biosynthesis.

Our analyses also revealed negative correlations between RLS and transcript abundance for all genes (negative correlation) involved in BCAA biosynthesis from pyruvate in long-lived strains (**Fig. S8, Table S2**). On the other hand, we did not observe any changes in Val, Leu and Ile abundance (**Table S2)**. This observation raises a possibility that intracellular homeostasis of these BCAAs might be regulated through other resources (e.g. extracellular import) [95].

The other metabolites that showed a significant correlation to RLS were lactic acid (LA), phenyl-lactic acid (PLA), tyrosine (Tyr) and aspartic acid (Asp) (**Table S2**). Although the synthesis of LA from pyruvate is well studied, the metabolic regulation and function of PLA, a product of the shikimate pathway, are less clear. Previously, it was found that yeast produces PLA through a nonspecific activity of lactate dehydrogenase from phenylpyruvate, a metabolite derived from chorismate in the shikimate pathway [96] **(Fig. S5)**. Interestingly, Tyr is also synthesized via the shikimate pathway (**Fig. S5**) and its abundance negatively correlates with RLS. The decreased *ARO2* (synthesizes chorismate from shikimate) expression might explain the decreased abundance of Tyr in long-lived strains. Similarly, one can expect a decreased Trp abundance, which is also synthesized through the shikimate pathway **(Fig. S5)**. However, our data revealed an increased level of Trp in long-lived strains. Therefore, we relate this observation to the decreased *BNA2* expression (see above).

Finally, since amino acid metabolism is directly related to glycolytic and/or TCA cycle intermediates **(Fig. 6)**, we analyzed differences in metabolites and genes involved in these central metabolic processes between short- and long-lived strains. Analysis of the data based on metabolomics and transcriptomics approaches suggested a decreased glycolytic rate and increased TCA cycle activity in long-lived strains **(Fig. 6)**. For example, we found that glycolytic genes such as *FBA1, TDH2, PGK1, ENO2* and *CDC19* were negatively correlated with RLS, while TCA cycle genes such as *CIT1, IDP1* and *KGD1* were positively correlated with RLS (**Table S2**). These observations are consistent with the pathway enrichment analysis revealing increased TCA cycle and oxidative phosphorylation in long-lived strains. To further examine this, we measured basal oxygen consumption rate (OCR) of wild yeast isolates and verified the increased respiration rate in long-lived strains. To determine if the observed pattern of median RLS variation can be partly explained by this increased mitochondrial function, we tested a potential relationship between OCR and median RLS. Our analysis revealed a significant positive correlation (*R*=0.28, *Padj*=0.016) between OCR and median RLS (**Fig. S9A, Table S5**). Furthermore, we tested whether the increased OCR can be simply explained by total mitochondrial copy number by analyzing protein abundance of mitochondrial marker protein Por1 by Western blots. We observed a similar abundance of Por1 across the strains, arguing against alteration in mitochondrial copy number in long-lived strains (**Fig. S9B**). Overall, this data suggests a possible role of mitochondrial function in lifespan variation across wild yeast isolates.

**Figure 6:**
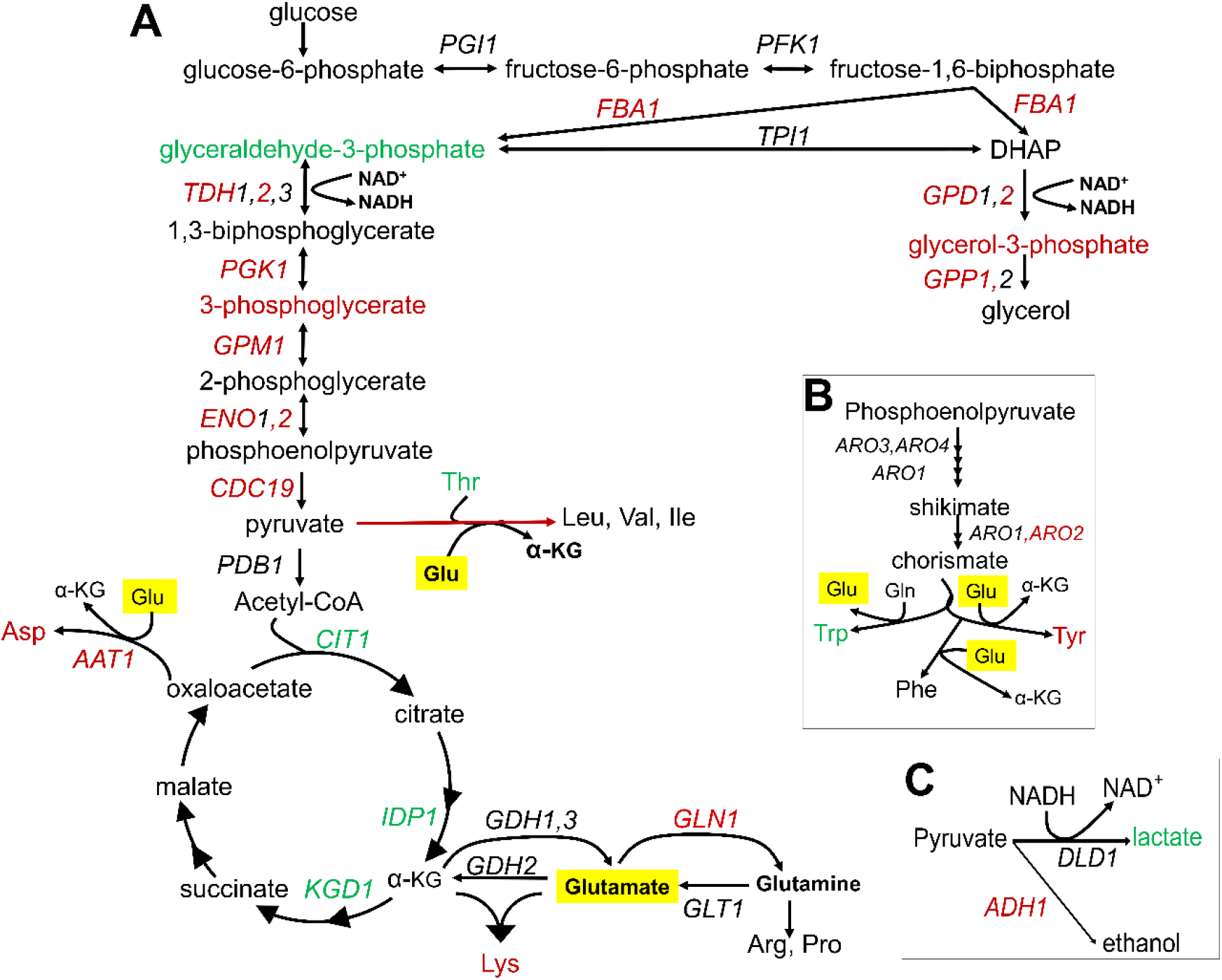
Summary of metabolic changes associated with RLS. **(A)** Summary depiction of genes and metabolites from the inter-connected glycolytic pathway, TCA cycle and amino acid metabolism that are found to be associated with RLS. Associated genes are colored in red (negatively associated with RLS) or green (positively associated with RLS). Depiction of **(B)** shikimate pathway and **(C)** lactate and ethanol biosynthetic pathways are shown with the same color code representation. Glutamate is highlighted in yellow.

In summary, our joint omics analyses revealed consistent changes associated with increased lifespan at both metabolome and transcriptome levels, pointing to decreased glycolytic activity and amino biosynthesis and increased mitochondrial activity (TCA cycle and mitochondrial respiration) even under conditions of excess fermentative carbon source (glucose). These findings suggest common changes responsible for modulating lifespan across a broad diversity of wild yeast isolates.

### Experimental testing of Glu and Trp metabolism in regulation of longevity

To further understand the molecular mechanisms that support the long life of yeast wild cells, we paid particular attention to the association between decreased Trp degradation (KYN pathway) and RLS. Our data highlight the importance of Trp metabolism in lifespan regulation in long-lived strains. We found that even though the Trp biosynthesis pathway (shikimate pathway) is suppressed, Trp levels were increased, possibly due to decreased transcript abundance of *BNA2*, which controls the first-rate limiting step in Trp degradation. This data suggest that Trp abundance itself might be important for longevity and that long-lived strains might compensate for decreased Trp biosynthesis by inhibiting Trp degradation. To test this idea, we examined the lifespan effect of increased *BNA2* dosage in three long- and short-lived strains. Consistent with the findings from transcriptomic data, the increased expression of *BNA2* caused a significant decrease in median RLS in two out of three long-lived strains tested and significantly decreased maximum RLS in all long-lived strains tested (**Fig. 7A**). The increased expression of *BNA2* in short-lived strains did not result in a consistent RLS pattern. Among the three short-lived strains tested, two decreased median and maximum RLS significantly, and caused a significant increase in median RLS in one strain **(Fig. 7B)**. Additional data are needed to fully clarify whether changes in Trp levels or some intermediate metabolites from KYN pathway such as QA are critical for the observed lifespan variation; however, our data support a role for Trp homeostasis in longevity.

**Figure 7.**
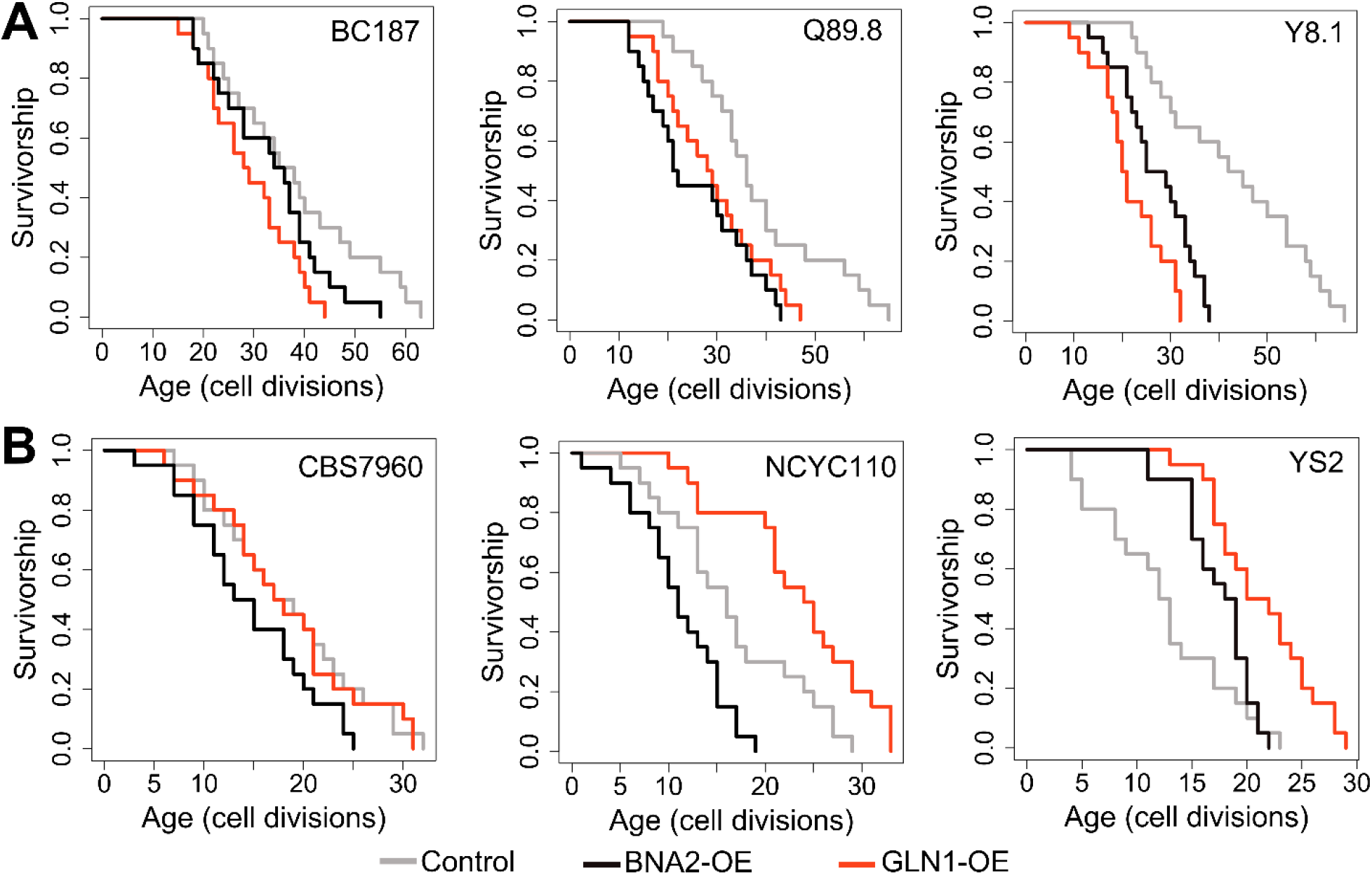
RLS effect of *GLN1* and *BNA2* overexpression in selected long- and short-lived strains. Lifespan curves for control (Gray), *BNA2* (Black), and *GLN1* (red) overexpression in (A) long and (B) short lived strains. Lifespan data and significance of lifespan changes can be found in Supplementary Table 1.

The observation of decreased glycolysis, increased SCFAs abundance and increased lactic acid synthesis from pyruvate in long-lived strains seems to disagree with the findings from transcriptome and the experimental work that suggest increased TCA cycle activity and respiration. While our findings from both metabolomics and transcriptomics data suggest a decreased substrate availability for the TCA cycle, increased TCA cycle activity may be fueled by alternative substrates. We hypothesize that compartment-specific glutamate (Glu) to alpha ketoglutarate (α-KG) flux, a reaction mainly controlled by NAD^+^ dependent mitochondrial Glu dehydrogenase, *GDH2* in mitochondria [97–99], might support increased TCA activity. In this case, spared Glu (due to decreased Glu-dependent amino acid biosynthesis) can support citrate synthesis to fuel TCA cycle via α-KG conversion. Along with decreased glycolysis and a decrease in Glu-dependent amino acid biosynthesis, compartment specific Glu to α-KG flux might be important for extended longevity in long-lived strains. In fact, our findings suggest that Glu utilization is limited in long-lived strains; however, the observation of no significant alteration in Glu abundance is consistent with the idea that long-lived strains may utilize Glu in some other pathway. In support of this model, we found that transcript abundance of *GLN1*, an enzyme responsible for synthesis of glutamine (Gln) from Glu (**Fig. 6**) in mitochondria, negatively correlates with lifespan (**Table S2**). To test the possibility that *GDH*-mediated Glu to α-KG flux is important for supporting the lifespan of long-lived strains, we overexpressed *GLN1* in three long- and three short-lived strains. Overexpression of *GLN1* is expected to decrease the Glu pool, and thus perhaps α-KG synthesis. We found that *GLN1* overexpression significantly decreased both median and maximum RLS of long-lived strains tested (Wilcoxon rank sum tests, p < 0.05) **(Fig. 7A)**. Among the three short-lived strains tested, two significantly increased median and maximum RLS, while the remaining strain showed no significant lifespan changes **(Fig. 7B)**. Thus, the data support the idea that the mitochondrial Glu pool may have a role in longevity across wild isolates. Although it needs additional experimental evidence, we think that increased NADH levels in long-lived strains might be due to increased NAD^+^-dependent *GDH2* activity, which catalyzes the conversion of Glu to α-KG in mitochondria.

Although initially yeast was considered as Krebs-negative (i.e. cannot utilize TCA cycle intermediates as carbon sources for growth) [100], later on α-KG was shown to be catabolized, under the condition of co-consumption with low glucose [101]. To investigate whether wild isolates can utilize α-KG as an alternative carbon source, we cultured them in the medium containing low glucose and α-KG, α-KG only and YP (yeast extract peptone without glucose) medium (**Fig. 8A, Fig. S10**). To our surprise, we found that many of these wild isolates showed weak growth even on the YP medium, which was not supplemented with any carbon source (**Fig. S10**). It is possible that some compounds in yeast extract may promote weak growth of these isolates, and we think that it might be α-KG. To prove this, we supplemented YP and YPD medium with α-KG (10g/l) and observed that many of the strains showed improved growth, further supporting utilization of α-KG for growth on the medium lacking glucose (**Fig. 8A, Fig. S10**).

**Figure 8:**
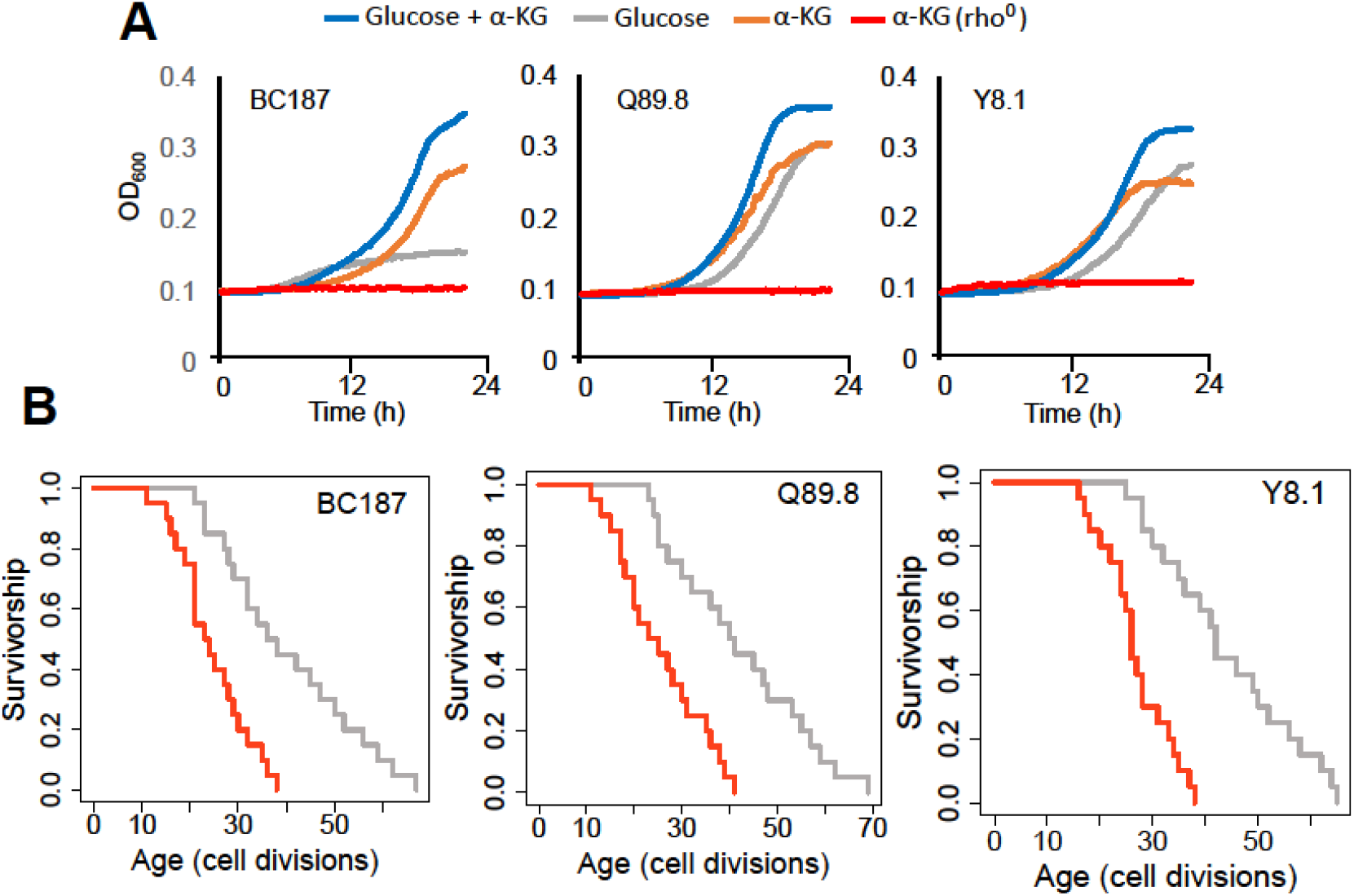
Growth properties of long-lived trains in medium supplemented with α-KG and effect of mtDNA elimination on α-KG utilization and RLS. **(A)** The growth of three long-lived strain were further supported with α-KG supplementation (10g/l). However, strains lost the ability of α-KG utilization upon mtDNA elimination (red). Growth data of OD_600_ measurement can be found in Supplementary Table 1. **(B)** Elimination of mtDNA significant reduced RLS in all three long lived strains. Lifespan data and significance of lifespan changes can be found in Supplementary Table 1.

Finally, to connect increased respiration, α-KG utilization and extended lifespan, we eliminated mitochondrial DNA (mtDNA, rho^0^) in three long-lived strains. We assayed their growth in medium supplanted with α-KG. We found that elimination of mtDNA in long-lived strains abolished their growth ability in the medium supplemented with α-KG as a sole carbon source (**Fig. 8A**). We further measured RLS of these rho^0^ isolates under 2% glucose conditions to understand whether blocking respiration would affect their lifespan. Our analysis revealed that the loss of mtDNA caused a significant reduction in RLS in all three strains tested (Wilcoxon rank sum tests, p < 0.05) (**Fig. 8B**). Overall, these data further support the idea that α-KG utilization and increased mitochondrial respiration are connected to each other and utilization of α-KG requires active mitochondria. Perhaps, under the conditions of decreased amino acid synthesis α-KG utilization could increase respiration which in turn may increase lifespan.

## DISCUSSION

The budding yeast has contributed significantly to our understanding of genetics and cell biology and has become an important model of aging, ever since Mortimer discovered the yeast RLS phenotype [13]. With the power of genetics and experimental tools, yeast has provided various clues for understanding the aging process in eukaryotes and yielded hypotheses that have been further tested in other organisms, including mammals [102, 103]. From this perspective, the natural isolates we analyze in the current study offer an excellent new model for yeast aging studies [18, 53–57], allowing us to leverage the enormous genetic variation found among natural isolates to study cellular processes that affect lifespan variation in nature, in a way not possible with the standard approach of deletion mutants in lab strain. In fact, these studies revealed several previously known, as well as novel, cellular processes and genetic factors that together determine replicative lifespan of yeast. For example, QTL analysis revealed a possible role of rDNA origin activation, nutrition sensing pathways and serine biosynthesis in modulation of replicative and chronological lifespan in wild yeast isolates. In addition, both initial and age associated increase in cell size found to be negatively correlated with RLS. In general, these studies pointed out diet-dependent metabolic regulations in lifespan regulation [18, 53–57].

In this study, we further advanced these findings by utilizing -omics approaches across highly diverse aging phenotypes. Our comparison of gene expression changes and longevity signatures across laboratory-adapted long-lived mutants and long-lived natural isolates identified many genes and pathways associated with longevity. However, further studies are needed to determine their individual and collective roles in lifespan variation. At the pathway level, the transcriptomic and the metabolomic data suggest that respiratory metabolism is important for longevity, as long-lived strains are characterized with increased TCA cycle and oxidative phosphorylation activities. This is also consistent with prior data that genetic induction of respiration in the PSY316 laboratory-adapted strain is sufficient to increase RLS [104]. In addition, we found that short-lived strains when grown with glucose as the primary carbon source (YPD) tend to achieve the largest lifespan gains when grown on glycerol (YPG) that induces a metabolic shift away from fermentation and toward respiration. In contrast, strains that are long-lived on YPD generally did not show a further RLS increase. Taken together, these findings suggest possible adaptive mechanisms under glucose conditions that suggest a metabolic shift from fermentation to respiration to increase mitochondrial metabolism in long-lived isolates. Accordingly, further increase in respiration by shifting the carbon source from glucose to glycerol was not beneficial in those long-lived strains.

In addition, our combined analyses of transcriptome and metabolome data pinpointed a regulation of interconnected amino acid biosynthetic pathways, which are down-regulated in long-lived strains. Amino acids are the building blocks of proteins, and it is known that individual supplementation or restriction of several different amino acids can exert both pro- and anti-longevity effects mainly through a well-studied target of rapamycin (TOR) pathway [105, 106]. For example, restriction of BCAAs was shown to increase both healthspan and longevity in mice [107, 108]. Jiang and colleagues first reported that reducing the amino acid content of the media can increase RLS in a short-lived laboratory yeast strain [109]. Similarly, Asp restriction [110] or treatment with the glutamine synthetase inhibitor methionine sulfoximine [111] can extend RLS by inhibiting TOR. Our data are consistent with the idea that decreased amino acid biosynthesis plays a role in a longer lifespan of wild isolates and that this is associated with decreased glycolytic activity and increased mitochondrial function.

Among the amino acids that are found to be associated with lifespan, Trp appears to be particularly relevant for lifespan regulation. Recently, a decrease in KYN metabolic pathway activity through RNAi knock-down of *TDO-2 (BNA2* ortholog) expression (knock-down) was found to robustly extend lifespan, [112] while complete knock-out of TDO-2 expression diminished the positive lifespan effect in *C. elegans* [113]. Increased KYN pathway activity and alteration in KYN pathway metabolites with age have also been observed in humans, suggesting a possible conserved role for this pathway in lifespan regulation [114, 115]. In addition, a study across 26 mammalian species found that species characterized by increased KYN pathway activity were shorter-lived [110]. In yeast, it was shown that deletion of *BNA2*, which encodes the protein that controls the first rate limiting step in Trp catabolism, decreased RLS, while increased *BNA2* dosage (overexpression) increased RLS in diploid laboratory WT cells [116]. We observed that transcript abundance of *BNA2* negatively correlated with lifespan across wild isolates. Consistent with this observation, we found that increased *BNA2* dosage caused a significant lifespan reduction in long-lived strains as well as short-lived strains. These data support the model that decreased KYN pathway activity is associated with increased lifespan across wild isolates. Due to decreased shikimate pathway activity, increased *BNA2* dosage possibly caused increased activation of the KYN pathway by increasing Trp degradation, which in turn resulted in decreased intracellular Trp pool for protein translation in long-lived strains. On the other hand, short-lived strains are already characterized with increased *BNA2* abundance and further increase in *BNA2* dosage might increase KYN pathway metabolic intermediates (e.g. kynurenine, QA) and result in further lifespan reduction. Perhaps, the direct way to test the role of Trp in lifespan regulation should be to analyze the effect of decreased expression of *BNA2* in short-lived strains, which will directly increase the abundance of Trp, in a similar fashion to that in long-lived strains. Although our data differ from the recently published report in which *BNA2* overexpression increased lifespan [116], it is possible that Trp metabolic regulation and KYN pathway activity might follow different metabolic and genetic trajectories across wild isolates in comparison to the laboratory adapted strain used in that study, which had been cultured on a medium with high glucose and abundant Trp over many generations.

The KYN pathway activity could also be related to NAD^+^ homeostasis since Trp degradation is the major route for NAD^+^ synthesis [117]. Accordingly, we hypothesized that the observed unchanged NAD^+^ abundance across yeast isolates might be explained by the increased activity of the downstream NAD^+^ salvage pathway **(Fig. S5)**. In fact, we found that the expression of the nicotinamidase gene, *PNC1*, in the salvage pathway positively correlates with lifespan **(Fig. 3)**. Nicotinamide (NAM) is a by-product generated during Sir2p-mediated deacetylation and can be taken up from the medium. The stress-induced nicotinamidase Pnc1p in yeast is responsible for the clearance of NAM by converting it to nicotinic acid (NA), which is a precursor for NAD^+^ biosynthesis via the salvage pathway [118, 119] **(Fig. S5)**. Increased expression of *PNC1* alone has been shown to modulate intracellular NAD^+^ homeostasis and to increase RLS [116, 117]. In addition to the hypothesis that the increased salvage pathway activity might compensate for NAD^+^ biosynthesis in long-lived strains with decreased KYN activity, our finding of increased lactate abundance in these strains could be interpreted as an alternative route for NAD^+^ regeneration. During lactic acid fermentation, two molecules of pyruvate are converted to two molecules of lactic acid. This reaction also supports oxidation of NADH to NAD^+^. Previously, it has been shown in both yeast and mammalian cells that when NAD^+^ demand is higher relative to ATP turnover, cells engage in anaerobic glycolysis, despite available oxygen [120]. Our data also suggest that a similar mechanism might have evolved to regulate NAD^+^ homeostasis in cells with decreased KYN pathway activity. Overexpression of *BNA2* might also interfere with these adaptive changes in long-lived strains, which in turn decreases lifespan. Our molecular identification of adaptive metabolic changes may prove useful in uncovering additional mechanisms regulating cellular NAD^+^ metabolism and their association with the aging process in future studies.

A potential connection between altered amino acid biosynthesis and the TCA cycle that may be particularly relevant for lifespan determination is Glu metabolism. Other than being a precursor in many amino acid biosynthetic pathways, Glu is an important carbon and nitrogen carrier, and can be catabolized to α-KG, an intermediate of the TCA cycle through a deamination reaction catalyzed by GDH2 as well as by other transaminases such as *BAT1, BAT2* and *ARO8* during amino acid biosynthesis. The movement of α-KG through the TCA cycle represent the major catabolic step for the production of nucleotides, lipids, and amino acids [121]. Here, we also showed that wild yeast isolates can use α-KG as an alternative carbon source for growth. We hypothesize that mitochondria specific Glu to α-KG conversion by GDH2 might be an important determinant of lifespan regulation. In fact, increasing utilization of the Glu pool towards Gln resulted in a significant decrease in lifespan in long-lived strains. Based on these data, both compartment specific Glu to α-KG conversion by GDH activity and utilization of α-KG for energetic and/or anabolic purposes might result in longer lifespan across wild isolates. Hence, our data suggest a possible mechanism that niche-specific nutrient depletion promotes halting the biosynthetic machinery (e.g. amino acid biosynthesis, glycolysis) and alleviates catabolic processes of alternative carbon sources to provide energy maintenance in long-lived strains by increasing respiration. Recently, α-KG emerged as a master regulator metabolite [122]. There have been many enzymes found to be regulated by α-KG, characterized as an epigenetic regulator, and identified as a regulator of lifespan in *C. elegans* [123] and mouse [124]. In *C. elegans*, α-KG was found to decrease ATP levels by blocking mitochondrial complex V activity, thereby reducing oxygen consumption. This effect was found to be mTOR-dependent [123]. Similarly, α-KG was found to extend lifespan in fruit fly by inhibiting mTOR and activating AMPK signaling [125]. In mid-aged mice, α-KG supplementation decreased systemic inflammatory cytokines leading to health and lifespan benefits [124]. More recently, an analysis of 178 genetically characterized inbred fly strains revealed α-KG-dependent lifespan regulation under dietary restricted conditions [126]. However, in yeast, the effect of α-KG supplementation on lifespan regulation has never been tested. It was shown that yeast can actively transport α-KG from medium to cytosol and into mitochondria [100, 101]. In addition, in contrast with the findings in *C. elegans*, α-KG supplementation was found to increase oxygen consumption in yeast [100]. Also, α-KG supplementation was shown to increase oxidative stress resistance in yeast [127]. Similarly, a dietary role of Glu in aging has been tested in different model organisms, including yeast. Initially, Glu restriction was found to increase yeast chronological lifespan (CLS) [128]; however, later on, it was shown that Glu supplementation also has a positive effect on CLS [110]. Similarly, in *C. elegans*, medium supplemented with a lower dose (1-5 mM) of Glu was found to extend lifespan [105]. However, the role and mechanisms of Glu metabolism in lifespan regulation are not well understood. Both α-KG and Glu are involved in epigenetic and redox regulations that all have been implicated in lifespan regulation [124, 129] and might also provide a mechanism for lifespan extension in long-lived strains. Furthermore, catalysis of Glu to α-KG also yields NH4, which has been shown to be involved in regulation of mTOR1 and mTOR2 signaling [130, 131] and lifespan. All these findings from different organisms suggest complicated mechanisms of beneficial effects of a-KG, which needs further investigation.

Overall, our research takes advantage of natural variation in yeast lifespan that has arisen in response to mutation, selection and genetic drift, and uses this variation to identify the potential causal roles that gene expression and metabolism play in shaping lifespan within the same species. Our data revealed a novel mechanism wherein different life history trajectories contribute to mitochondrial metabolism. Hence, the TCA cycle represents a central metabolic hub to provide metabolites to meet the demands of proliferation and other cellular processes. With respect to this, modification of TCA metabolic fluxes and metabolite levels in response to environmental pressures might therefore account for cellular adaptation and plasticity in the changing environment which might also affect lifespan of these wild isolates. We further provide molecular insights into the unique metabolic adaptation involving linked pathways, involving in Glu and a-KG metabolisms in regulation of mitochondrial function and their possible association with lifespan variation. Further understanding of how gene-environment interactions modulates genes and pathways associated with longevity may open new therapeutic applications to slow aging and delay the onset of age-related diseases through diet, lifestyle, or pharmacological interventions. In future studies, it might yield important information to investigate the role of α- KG metabolism in amino acid and caloric restricted lifespan regulation.

## Materials and Methods

### Yeast strains and growth conditions

Many of the diploid wild isolates of *S. cerevisiae* and *S. paradoxus* (68 isolates) were obtained from the Sanger Institute [49] and the remaining 8 isolates of *S. cerevisiae* were gifted by Justin Fay from Washington University [48]. Detailed information about strains used in this study is in Supplementary Table 1. The diploid laboratory WT strain BY4743 was purchased from the American Type Culture Collection. For testing the growth effect, strains were cultured overnight in a 96–well plate incubator at 30°C in YPD medium. Next day, 1μl from overnight culture was transferred to the YP, YP + α-KG (10g/l) or YPD (0.02% glucose) + α-KG and growth was monitored in 96 well plate using Epoch2 (BioTek, Winooski, VT, USA) kinetic growth analyzer by analyzing optical density of OD_600_. For expression of genes of interest, we used modified p426GPD high copy plasmid by inserting a hygromycin (HYG) cassette along with its promoter and terminator at the *XbaI* restriction site. HYG cassette was amplified from pGAD32 plasmid with PCR. Using modified p426GPD, we inserted *GLN1* and *BNA2* gene cassettes individually at the *BamHI/XhoI* restriction sites for overexpression. Yeast transformation was performed using standard lithium acetate method. Growth rates were determined using a BioScreen-C instrument (Bioscreen C MBR, Piscataway, NJ, USA) by the analysis of optical density in the OD_600_ range, and doubling times were calculated with an R script by analyzing fitting spline function from growth curve slopes [132]. The maximum slope of the spline fit was used as an estimate for the growth rate and doubling time for each evolved line, in combination with the YODA software package [133]. Finally, mtDNA was eliminated by culturing cells in YPD medium, supplemented with 10 μg/ml and ethidium bromide (EtBr). Briefly, logarithmically growing cells (OD_600_=0.5) were incubated at room temperature with agitation for approximately 24 h. Following a second and third treatment with the same concentration of EtBr for 24 h, the cells were diluted (1:100) in water and plated on YPD to obtain single colonies. After then, several individual colonies were selected for testing their growth ability on YPG plates. Colonies, which were unable to grow on YPG were selected as rho^0^.

### Replicative lifespan assay

RLS was determined using a modification of our previously published protocol [134]. Yeast cell cultures for each strain were freshly started from frozen stocks on YPD plates and grown for 2 days at 30 °C prior to dissections. Several colonies were streaked onto new YPD with 2% glucose, YPD with 0.05% glucose or YPG plates with 3% glycerol using pipette tips. After overnight growth, ~100 dividing cells were lined up. After the first division, newborn daughter cells were chosen for RLS assays using a dissection microscope. For each natural isolate, at least two independent assays were performed using at least sets of 20 cells for each assay. Each assay included 20–80 mother cells of BY4743 strain as well, which was used in every experiment as a technical control. For RLS analysis of wild isolates harboring expression plasmids, individual colonies were picked up from selection medium (HYG) and YPD medium supplemented with 200 microgram/mL HYG were used for RLS determination of these cells. Survival analysis and Gompertz modeling was performed using the survival and flexsurv packages in R, respectively.

### Measurement of basal oxygen consumption rate and western blot analysis

To investigate metabolic respiration differences across wild isolates OCR, (pmol/min) was measured using a Seahorse XFe96 analyzer (Agilent, Santa Clara, CA, USA). Cells grown overnight in YPD were diluted to OD600=0.01 in the morning, and cells were grown to reach the OD600=0.25-0.5. Then, cell culture was diluted to OD600=0.02 in YPD and placed in a XFe96 cell culture plate coated with 15 microliter 0.01% poly-L-lysine and attached to the plate according to the previously published protocol [135]. Basal OCR was measured for 5 cycles at 30 °C. To examine the expression of mitochondrial proteins, western blotting was carried out with antibodies against mitochondrial outer membrane protein Por1 (Abcam, Cambridge, MA, USA, cat:ab110326). For each strain, 10 mL logarithmically growing cells were collected and proteins were isolated according to previously published protocol [18]. The membranes were stripped and developed with antibodies against phosphoglycerate kinase (Pgk1; Life Technologies, Grand Island, NY, USA, cat: 459250) as an internal loading control.

### RNA-sequencing and data analysis

Three independent cultures for each strain were collected at the OD_600_=0.4 on YPD medium to isolate RNA from each culture using Quick-RNA 96 Kit from Zymo Research (Cat. number: R1053). To prepare RNA-seq libraries, Illumina TruSeq RNA library preparation kits were used according to the user manual, and RNA-seq libraries were loaded on Illumina HiSeq 4000 platform to produce 150 bp paired-end sequences. After quality control and adapter removal, the STAR software package [136] was used to map the reads against a pseudo reference genome of each strains, in which we replaced identified nucleotide changes in the S288c reference genome. Read alignment rate for transcriptome data against pseudo genome varied between 92-97% across *S. cerevisiae* strains and 93-99% across *S. paradoxus* strains (**Fig. S11**). Read counts per gene were calculated using feature Counts [137]. To filter out genes with low numbers of reads, we used filterByExpr function from the edgeR package and resulted in an expression set of 5,376 genes across replicates of wild isolates.

### Metabolite profiling and data analysis

A portion of the cell pellet collected for RNA-seq analyses was also used for targeted metabolite profiling using liquid chromatography-mass spectrometry (LC-MS). 1 ml of MeOH:H_2_O mixture (8:2, v/v) was added to the samples, swirled at 550 rpm on a mixer for 5 minutes and then transferred to an Eppendorf tube, they were sonicated in an ice bath for 10 min, centrifuged at 4°C at 14,000 rpm for 15 min, and 600 μl of supernatant was collected into a new tube and dried in a vacuum centrifuge at 30 °C for 2.5 hrs. Samples were reconstituted in 1 mL and injected into a chromatography system consisting of a dual injection valve setup allowing injections onto two different LC columns with each column dedicated to an ESI polarity. 5 μL were injected on the positive mode column and 10 μL on the negative side column. The columns were a matched pair from the same production lot number and were both a Waters BEH amide column (2.1 x 150 mm). Auto sampler was maintained at 4 °C and column oven was set to 40 °C. Solvent A (95% H_2_O, 3% acetonitrile, 2% methanol, 0.2% acetic acid with 10 mM ammonium acetate and 5 μM medronic acid and Solvent B (5% H_2_O, 93% acetonitrile, 2% methanol, 0.2% acetic acid with 10 mM ammonium acetate 5 μM medronic acid) were used for sample loading. After completion of the 18-minute gradient, injection on the opposite column was initiated and the inactive column was allowed to equilibrate at starting gradient conditions. A set of QC injections for both instrument and sample QC were run at the beginning and end of the sample run. Data was integrated by Multiquant 3.0.2 software. Peaks were selected based on peak shape, a signal-to-noise of 10 or better and retention times consistent with previously run standards and sample sets. Analysis of the dataset was performed using R (version 3.6.0). All the metabolites with ≥ 40% missingness were excluded, and a total of 166 metabolites were included in the imputation step. We imputed the remaining missing values using the K-nearest neighbors imputation method implemented in the R impute package. The log2-transformed abundance was Cyclic LOESS normalized prior to imputation.

### Principal component analysis (PCA)

Principal component analysis was performed on preprocessed data (e.g. normalized and imputed log2 abundance of the metabolomic data, and the log2-counts per million (CPM) values of the filtered and TMM normalized RNAseq data) using the R prcomp function. To identify the underlying pathways, the factors in each of the first three principal components were ranked by their contributions, and pathway enrichment analysis was performed on the top 500 transcripts using Network Analyst [138] and on the top 40 metabolites using MetaboAnalyst [89] platforms.

### Phylogenetic regression by generalized least squares

R packages ‘nmle’ and ‘phylolm’ were used to perform phylogenetic regression by generalized least squares method to identify RLS association of transcripts and metabolites [18]. We tested four models of trait evolution: (i) complete absence of phylogenetic relationship (‘Null’); (ii) Brownian Motion model (‘BM’); (iii) BM transformed by Pagel’s lambda (‘Lambda’); and (iv) Ornstein–Uhlenbeck model (‘OU’). The parameters for Lambda and OU models were estimated simultaneously with the coefficients using maximum likelihood. The best-fit model was selected based on maximum likelihood. Strength of correlation was based on the p-value of regression slope. To confirm robustness of results, regression was performed by leaving out each strain, one at a time, and computing *P* values using the remaining strains.

### Gene expression signature associated with RLS across deletion strains

Gene expression data on deletion mutants was obtained from GSE45115, GSE42527 and GSE42526 [86]. The corresponding RLS lifespan data for mutant strains was from [85]. Based on the raw data from the number of replicates, we calculated median, mean and maximum RLS, together with corresponding standard errors (SE) for each deletion strain. In total, this resulted in 1,376 deletion strains, for which both RLS and gene expression data were available. logFC of individual genes corresponding to each mutant strain compared to control samples were used for subsequent analysis.

To identify genes associated with RLS across KO strains linear models in limma were used [139]. We found genes associated with median, mean and maximum RLS both in linear and logarithmic scale, and BH adjustment was performed to account for multiple hypotheses [140]. Genes with adjusted p-value < 0.05 were considered significant. To determine statistical significance of the overlap between genes associated with different metrics of RLS, we performed Fisher exact test separately for up- and downregulated genes, considering 6,170 genes as background.

### Comparison between signatures of RLS across deletion and natural strains

To compare gene expression signatures associated with different metrics of RLS across deletion and natural *Saccharomyces* strains, we calculated Spearman correlation coefficients between corresponding gene expression slope coefficients in a pairwise manner. Clustering of the Spearman correlation matrix was performed with the complete hierarchical approach.

To increase the signal within the correlation matrix, the union of top 1000 statistically significant genes from each of the two signatures in a pair was used to calculate Spearman correlation coefficient. To get an optimal gene number for removal of noise, we looked at how the total number of significantly correlated pairs of signatures depended on the number of genes used to calculate the correlation coefficient. As a threshold for significance, we considered BH adjusted p-value < 0.05 and Spearman correlation coefficient > 0.1.

To determine statistical significance of the overlap between transcripts associated with different metrics of RLS across deletion and natural strains, we performed Fisher exact test, considering 4,712 genes as background. To identify genes whose deletions are associated with longer or shorter lifespan in *S. cerevisiae* strains, we compared the distribution of RLS across samples corresponding to certain deletion strains with the distribution of median RLS across all measured deletion strains. For that we used Mann-Whitney test. Genes with BH adjusted p-value < 0.05 were considered significant. Overlap of these genes with lifespan-associated genes across natural strains was assessed with the Fisher exact test with BH adjusted p-value threshold of <0.05.

### Functional enrichment analysis

For the identification of functions enriched by genes associated with RLS across deletion and natural strains, we performed gene set enrichment analysis (GSEA) [141] on a ranked list of genes based on log_10_(p-value) corrected by the sign of regulation, calculated as:

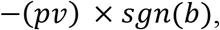

where *pv* and *b* are p-value and slope of expression of a certain gene, respectively, and *sgn* is signum function (is equal to 1, −1 and 0 if value is positive, negative and equal to 0, respectively). REACTOME, KEGG and GO biological process (BP) from Molecular Signature Database (MSigDB) were used as gene sets for GSEA [142]. We utilized the fgsea package in R for GSEA analysis. Adjusted p-value cutoff of 0.1 was used to select statistically significant functions.

We visualized several manually chosen statistically significant functions with a heatmap colored based on normalized enrichment score (NES). Clustering of functions has been performed with hierarchical complete approach and Euclidean distance. Combined integrative analysis of transcriptomics and metabolomics data for pathway analysis was performed by using joint-pathway analysis option in MetabolAnalyst 5.0 [89].

## Additional files Supplementary files

- Supplementary File. File contains supplementary figures and corresponding figure legends.
- Supplementary Table 1. Strain list, used in this study and their replicative lifespan along with doubling time are listed. This file also includes replicative lifespan of the laboratory KO strains, analyzed in this study.
- Supplementary Table 2. File includes raw and normalized counts from RNA-Seq. Normalized metabolite count is also included. Results from PGLS analysis from both datasets along with statistical values are listed as well.
- Supplementary Table 3. PCA loadings for normalized RNA-seq and metabolomics data sets.
- Supplementary Table 4. List of yeast longevity genes from GenAge database. Identified set of CR-related genes from our analysis is also listed.
- Supplementary Table 5. Values for oxygen consumption rate (OCR) measurement across the strains.
- Transparent reporting form

## Data availability

All data generated or analyzed during this study are included in the manuscript and supporting files.

## Author contributions

Alaattin Kaya, Conceptualization, Resources, Data curation, RLS analysis, Investigation, Visualization, Methodology, Writing original draft, Writing, review and editing, RNA-seq analysis, Project administration; Cheryl Zi Jin Phua, PGLS analysis, Visualization, Writing original draft; Mitchell Lee, Data curation, RLS and doubling time analysis, Visualization, Methodology, Writing original draft, Writing review and editing; Lu Wang, RNA-seq and Metabolomics data analysis, Visualization, Writing original draft; Alexander Tyshkovskiy, Data curation, Comparison of data between Wild isolates and laboratory KO strains, Visualization, Methodology, Writing original draft; Siming Ma, Methodology, PGLS analysis, Writing original draft; Benjamin Barre, Methodology, Writing and editing; Weiqiang Liu, Visualization, Methodology, RNA-seq data analysis; Benjamin Harrison, Sample preparation for metabolomics studies; Methodology; Xiaqing Zhao, Analysis of doubling time data; Xuming Zhou, Methodology, RNA-seq analysis; Brian M. Wasko, RLS analysis; Theo K. Bammler, Data curation, Methodology; Daniel E. L. Promislow, Resources, Data curation, Methodology, Writing original draft; Matt Kaeberlein, Resources, Data curation, RLS analysis, Writing original draft, Writing review and editing; Vadim N. Gladyshev, Conceptualization, Resources, Investigation, Methodology, Writing original draft, Writing, review and editing, Project administration.

