## Supplementary File for "Evolution of Natural Lifespan Variation and Molecular Strategies of Extended Lifespan"

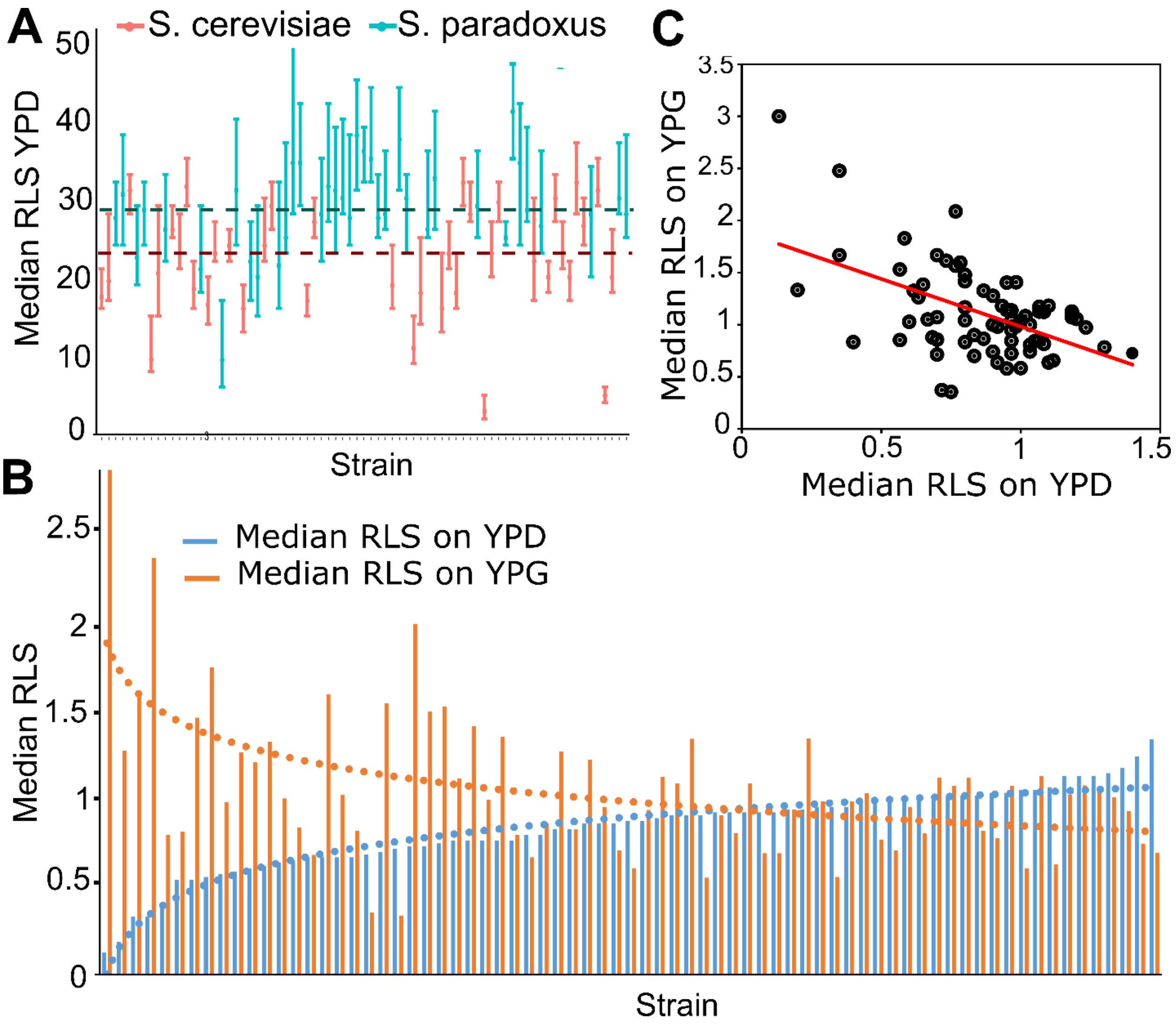


**Supplementary Figure 1: RLS phenotype across wild isolates.** A) Median RLS (Error bars are the 95%CI of the median lifespan) distribution across *S. cerevisiae* and *S. paradoxus* isolates grown in YPD. Dashed lines represent average median RLS of *S. cerevisiae* (red) and *S. paradoxus* strains (turquoise). **(B)** Median RLS changes in YPD (glucose-blue) and YPG (glycerol-orange) across wild isolates normalized against laboratory diploid WT strain, BY4743 (median RLS X strain / median RLS BY4743). Short-lived strains grown in YPD tend to have longer lifespan under YPG conditions, whereas long-lived strains grown in YPD tend to have shorter median RLS in YPG **(C)** Significant negative correlation between median RLS YPD and median RLS YPG. (Corr. coefficient = - 0.51, $P < 0.0001$).

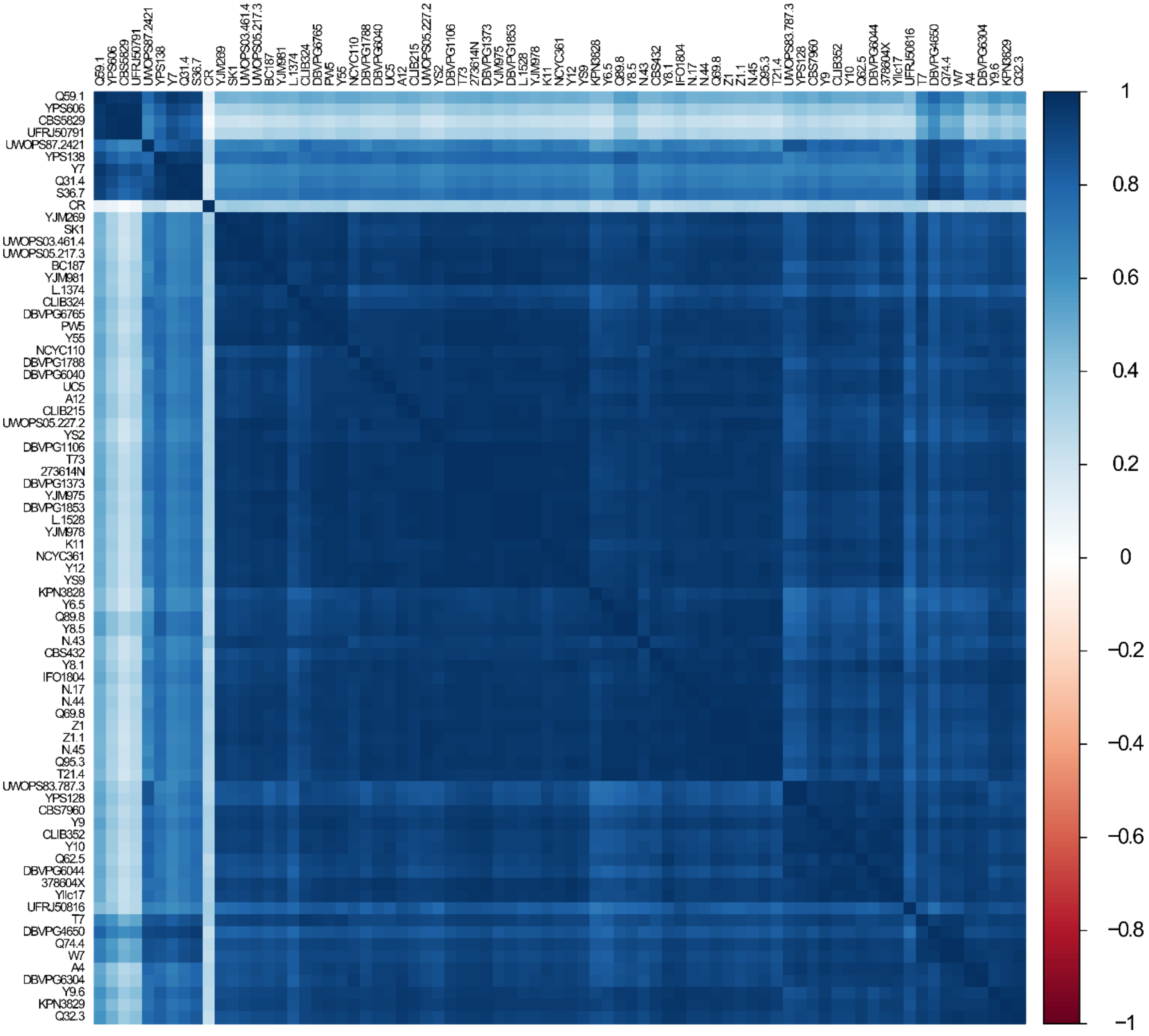


**Supplementary Figure 2: Correlation of genome wide transcript levels across wild isolates.** Heat map shows pairwise, 1-Pearson correlation matrix of gene expression data across wild isolates. Caloric restriction (CR) data was obtained from NCBI GEO, (SRA: SRX403444) and placed into the analysis.

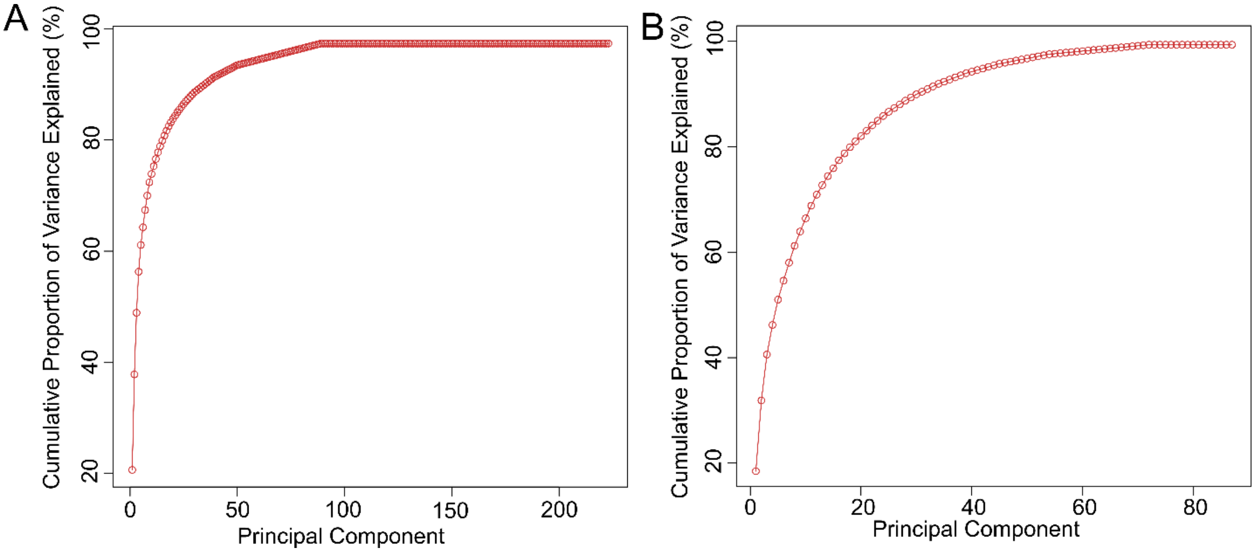


**Supplementary Figure 3: Principal component analysis.** Graphs show cumulative percentage of variance explained by Principal Components for (A) genes, (B) metabolites.

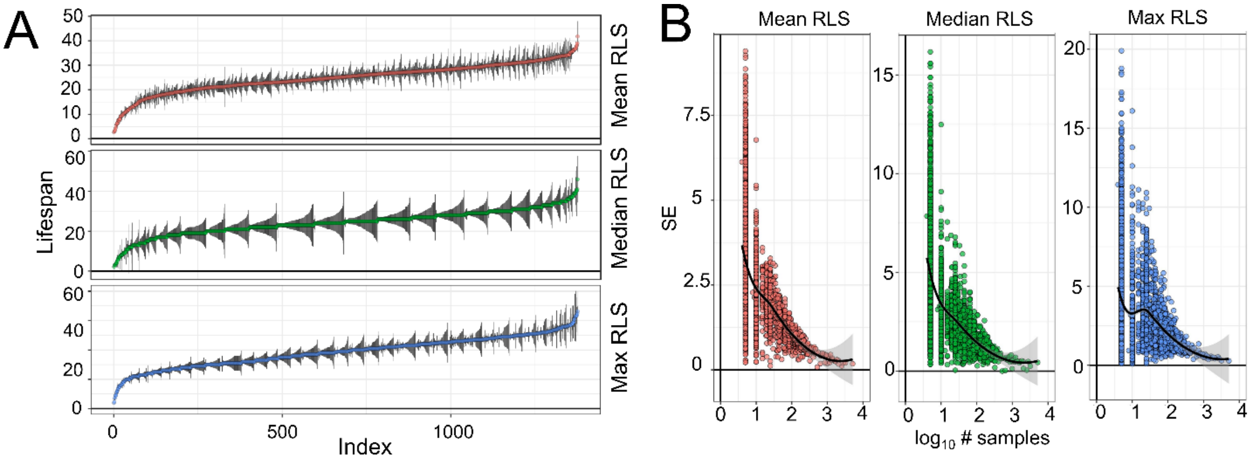


**Supplementary Figure 4: RLS phenotype of yeast knock-out strains.** A) Distribution of mean, median and maximum RLS across deletion strains with measured gene expression profile. RLS are sorted from the smallest to the greatest value. Black lines represent standard errors of the RLS estimates for corresponding strains. B) Dependence of standard error of RLS estimate on the number of strains used for evaluation of deletion mutant lifespan. Raw RLS values are provided in Supplementary Table 1.

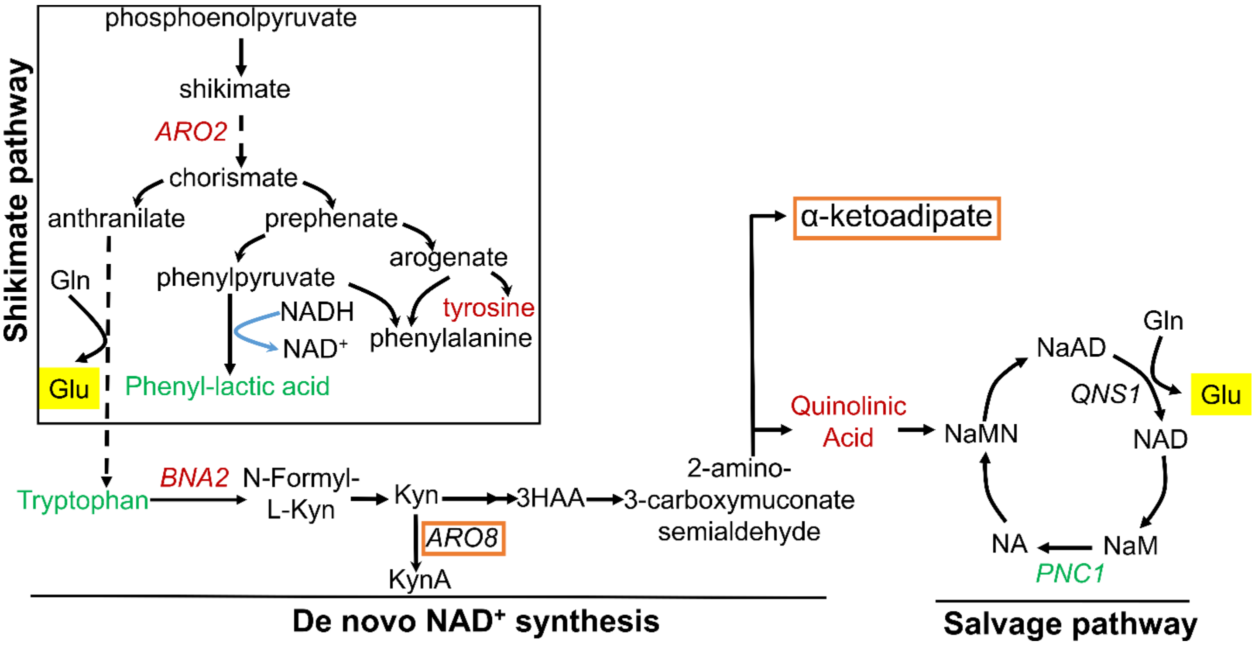


**Supplementary Figure 5: Genes and metabolites from shikimate, kynurenine and salvage pathways associated with RLS.** Genes and metabolites from each pathway that are found to be associated with RLS are colored with red (negatively associated with RLS) or green (positively associated with RLS). Genes and metabolites that are also known to be involved in lysine metabolism are highlighted in orange box, and glutamate is highlighted in yellow.

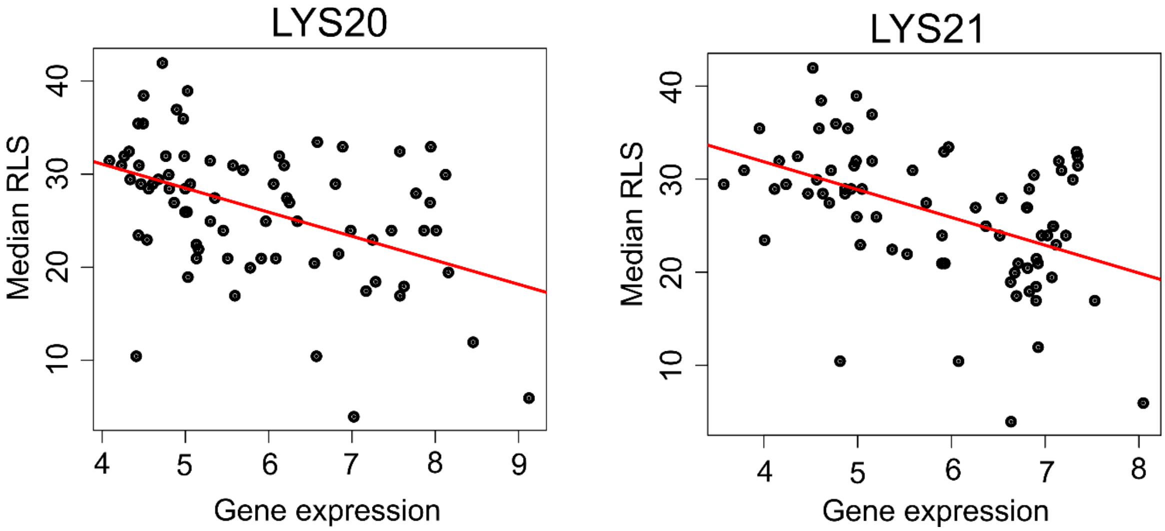


**Supplementary Figure 6: Correlation of LYS20 and LYS21 genes with median RLS.** Gene expression level (log2-cpm) of *LYS20* and *LYS21* negatively correlates with median RLS. Regression slope P values can be found in Supplementary Table 2.

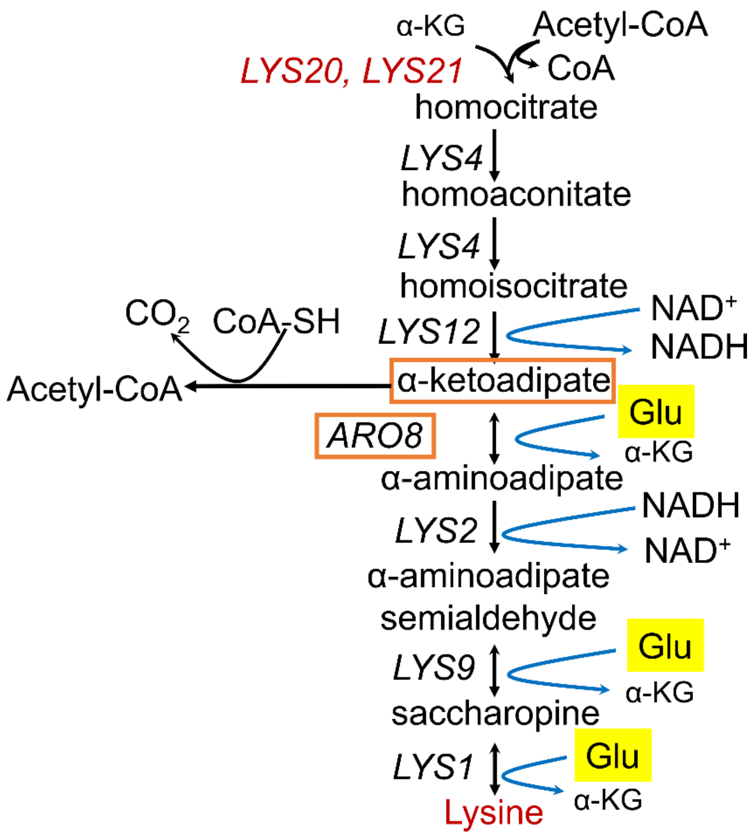


**Supplementary Figure 7: Lysine biosynthesis and RLS.** Genes and metabolites from the lysine biosynthetic pathway that are found to be negatively associated with RLS are colored in red. Genes and metabolites that are also known to be involved in tryptophan metabolism are highlighted with orange boxes, and glutamate is highlighted in yellow.

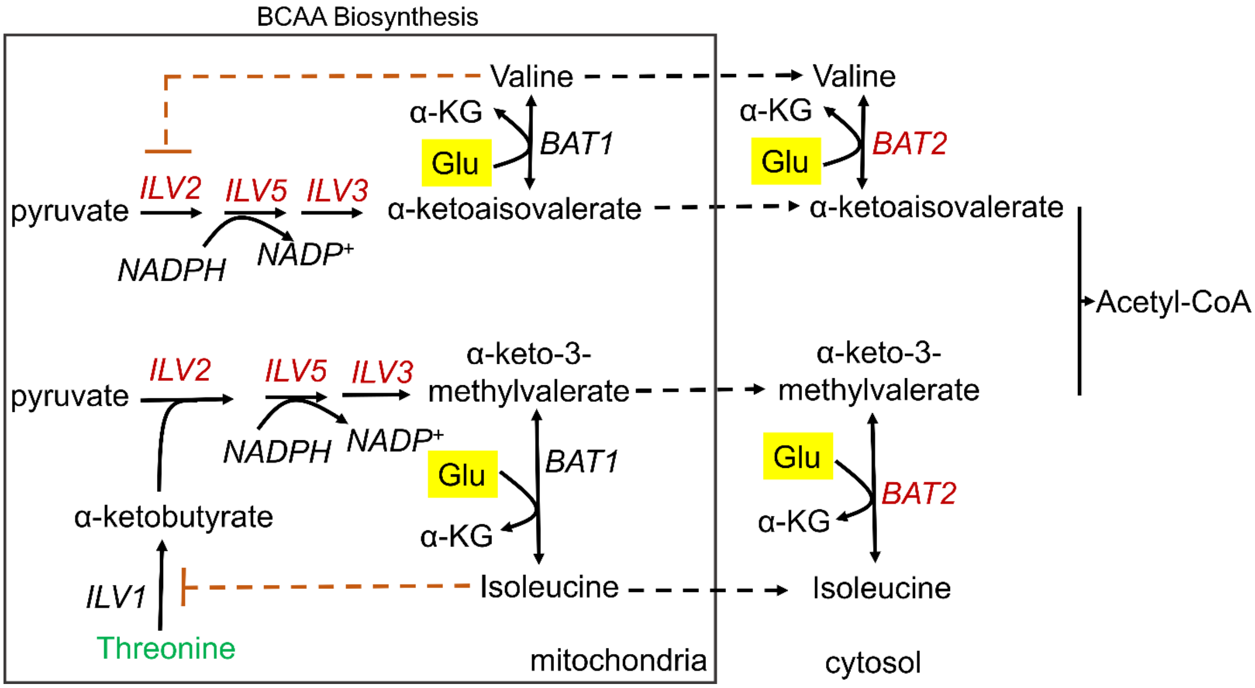


**Supplementary Figure 8: Genes from the BCAA metabolic pathway are negatively associated with RLS.** Genes involved in BCAA biosynthesis that are found to be negatively associated with RLS are colored in red. Threonine (positively associated with RLS) catabolism, is shown in green, and also contributes to the BCAA biosynthesis. Glutamate is highlighted in yellow.

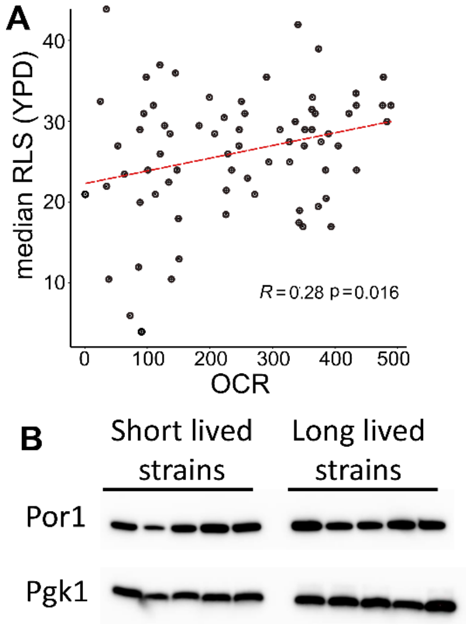


**Supplementary Figure 9: Association of mitochondrial respiration with median RLS. (A)** Spearman correlation between median RLS and OCR (pmol/min). **(B)** Western blot analysis for mitochondrial protein abundance across the strain. **S**imilar expression of mitochondrial porin Por1 (voltage-dependent anion channel) was verified with Western blotting. Pgk1 is used as internal loading control.

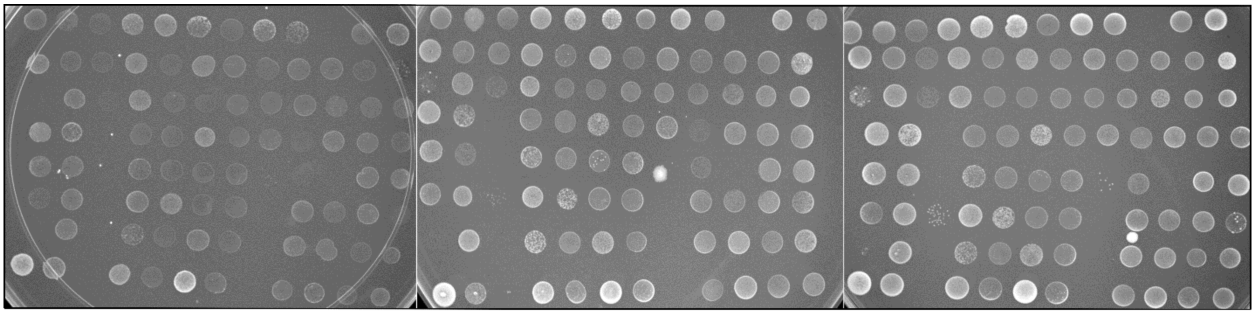


**Supplementary Figure 10. Effect of α-KG on growth.** Spot assay for wild isolates on medium containing YP (yeast extract and peptone), YP supplemented with α-KG (10g/l), and YP supplemented with both α-KG (10g/l) and glucose (0.02%). Each spot represents one of the strain. Plates were incubated at 30 $^{0}$C and pictured after 3 days.

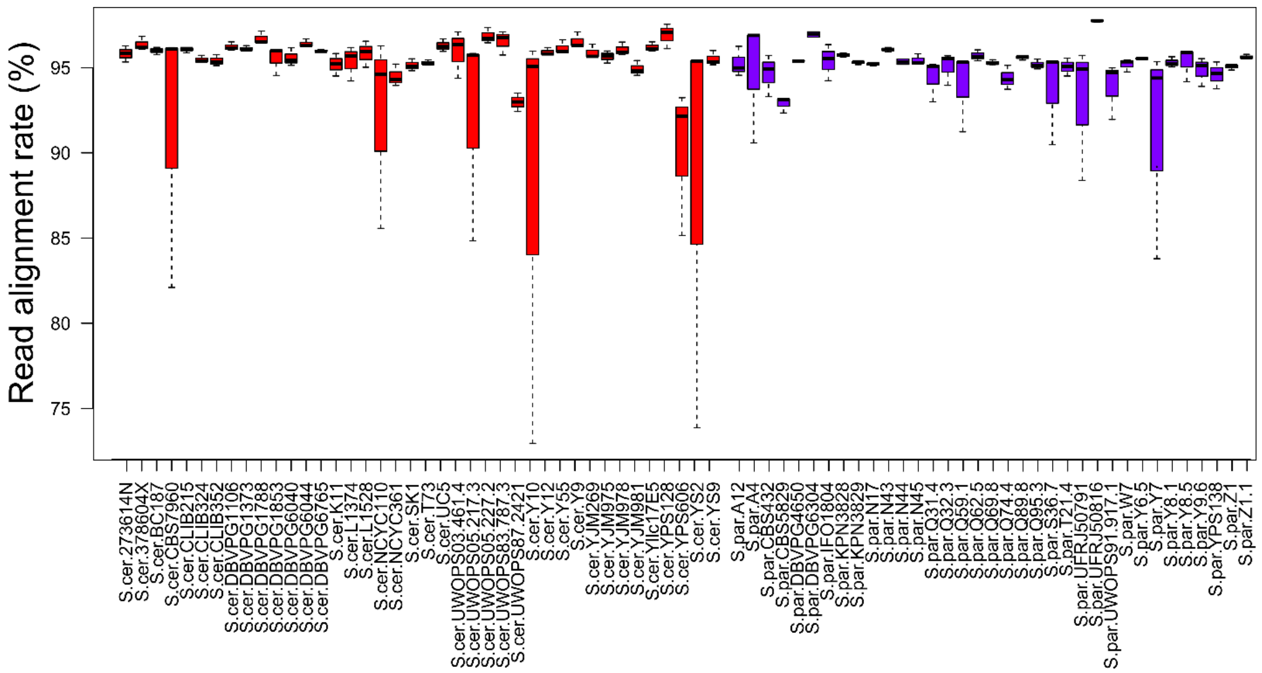


**Supplementary Figure 11. Read alignment rate for RNA-seq data.** RNA-seq reads were mapped to the pseudo genome using STAR aligner software and mean value of alignment rate (3 replicates per strain) from STAR outputs are shown here. Error bars show standard error of mean. Red bars indicate alignment rates for *S. cerevisiae* strains and purple bars indicate alignment rate for *S. paradoxus* strains.
